# Chronic abolition of evoked vesicle release from layer 5 projection neurons disrupts the laminar distribution of parvalbumin interneurons in the adult cortex

**DOI:** 10.1101/2024.05.06.592829

**Authors:** Florina P Szabó, Veronika Sigutova, Anna M Schneider, Anna Hoerder-Suabedissen, Zoltán Molnár

## Abstract

Establishing precisely built neuronal networks during cortical development requires appropriate proportions of glutamatergic and GABAergic neurons. Developmental disturbances in pyramidal neuron activity can impede the development of GABAergic neurons with long-lasting effects on inhibitory networks. However, the role of deep-layer pyramidal neurons in instructing the development and distribution of GABAergic neurons remains unknown. To unravel the role of deep-layer pyramidal neuron activity in orchestrating the spatial and laminar organisation of parvalbumin neurons, we selectively manipulated the activity of projection neurons in layer 5 of the cortex. By ablating SNAP25 from subsets of glutamatergic L5 projection neurons across the cortical mantle, we abolished Ca^2+-^dependent vesicle release from Rbp4-Cre+ pyramidal neurons. We explored the *local* (location of the cell bodies) and the *global* (subcortical projection sites) effects of chronically silencing cortical L5 neurons on parvalbumin interneurons. We found that the chronic cessation of vesicle release from L5 projection neurons left the density, distribution, and developmental trajectory of cortical and subcortical PV neurons intact during the second and third postnatal week; however, it resulted in the reorganisation of the laminar distribution of cortical PV neurons in layer 4 of S1 in the adult cortex. The abolition of evoked vesicle release from L5 also affected the perineuronal nets in the adult motor cortex and revealed a significant decrease in the density of VVA+ and PV-VVA+ cells in L5 of M1 at 3 months of age. The alterations in the laminar arrangement of VVA+ neurons may imply that Ca^2+-^ dependent synaptic transmission from L5 may control PNN density in adult networks. We also discovered that the absence of L5 activity only had a transient effect on the morphology of striatal PV neurons. The correlation between PV and VVA neurons was contingent on brain regions and cortical layers, therefore the link between the perineuronal net and PV neurons is significantly more intricate than previously believed. The present study will aid our understanding of the bidirectional relationship between deep-layer pyramidal cells and GABAergic neurons while uncovering the long-term effects of chronically disrupting pyramidal neurons on inhibitory networks.

## Introduction

The perplexing complexity of cortical circuits arises from the morphologically and functionally distinct cell types which are represented across the six layers of the neocortex and proven to have unique input and output connections. These intricately designed networks in the cerebral cortex rely on the precise ratio of excitatory pyramidal neurons and inhibitory GABAergic neurons that provide domain-specific innervation to distinct subclasses of pyramidal cells defined by their areal, laminar, and long-range projection targets (Kawaguchi and Kubota, 1997; Somogyi et al., 1998; Kepecs and Fishell, 2014; Markram et al., 2004; Tremblay et al., 2016). Molecularly distinct subtypes of inhibitory interneurons play a pivotal role in shaping circuit functions, gating information flow to the cerebral cortex, and synchronising the activity of neural assemblies in oscillatory neuronal networks (McBain and Fisahn, 2001; Somogyi and Klausberger, 2004; Gupta et al., 2000). In rodents, the GABAergic and glutamatergic neurons are generated in different sectors of the neuroepithelium under different combinatorial transcription codes (Butt et al., 2008; Marín and Rubenstein; 2011; Price et al., 1992; Sultan et al., 2013, Lodato and Arlotta, 2015). Disrupting the transcriptional identity of intratelencephalic (IT) projection neurons by knocking out the transcription factor Satb2 from pyramidal neurons not only converted the IT-type excitatory neurons to pyramidal-tract (PT) type neurons, but also caused aberrations in the cortical lamination of caudal ganglionic eminence (CGE)-derived interneurons (Wester et al., 2019). Similarly, loss of the transcription factor Fezf2 defining the fate of subcerebral projection neurons led to an aberrant laminar distribution of parvalbumin and somatostatin interneurons along with severe defects in GABAergic inhibition (Lodato et al., 2011). These studies suggest that both the transcriptional and the laminar identity of pyramidal neurons form an essential part of the cortical cytoarchitecture and may instruct the integration of GABAergic neurons into cortical circuits.

Although several subtypes of GABAergic neurons have long been studied for their ability to regulate pyramidal cell activity, less attention has been paid to the role that pyramidal neurons play in the establishment of inhibitory circuits. However, studies have emerged where perturbations in pyramidal cell function or local ablation of pyramidal cells have directly affected the number, survival, and synaptic connectivity of GABAergic interneurons (Lodato et al., 2011; Wong et al., 2018, 2022; Duan et al., 2020; Sreenivasan et al., 2022). Interneuron survival in the nascent cerebral cortex is dependent on the neuronal activity of pyramidal cells that rescue them from undergoing apoptosis by strengthening their connections to GABAergic cells before the programmed cell death (Denaxa et al., 2018; Wong et al., 2018, 2022; Sreenivasan et al., 2022; Priya et al., 2018). The interdependency of neuronal signalling in glutamatergic pyramidal neurons and the survival of GABAergic neurons points to the role of pyramidal neuron activity in the assembly of inhibitory cortical circuits. The intrinsically programmed mechanism of interneuron apoptosis itself can be modulated by chemogenetically activating pyramidal neurons and/or inhibiting PTEN signalling in pyramidal cells during the critical window of interneuron cell death in mice (Southwell et al., 2012; Wong et al., 2018). When pyramidal neurons are temporarily activated chemogenetically, parvalbumin and somatostatin interneuron density rises; conversely, when pyramidal neuron activity is manipulated during crucial stages of postnatal development, it decreases with transient inhibition. (Wong et al., 2018). While these studies have primarily focused on the regulatory role of pyramidal neurons on the survival of cortical GABAergic interneurons, new evidence suggests that PV interneuron survival in subcortical projection regions of cortical pyramidal neurons depends on the presence of glutamatergic inputs. When excitatory inputs from corticostriatal layer 5 projection neurons were absent, the number of striatal interneurons declined (Sreenivasan et al., 2022). This further supports the role of glutamatergic transmission and cortical control over the cortical and subcortical subtypes of PV interneurons. Ablation of the vesicular glutamate transporters of Vglut1 and Vglut2 to eliminate glutamate release from pyramidal neurons has been shown to have a differential impact on the number of GABAergic interneurons. While the abolition of excitatory transmission from pyramidal neurons has caused a marked decrease in the density of neurogliaform and basket cells, it has not altered the density of bipolar neurons (Wong et al., 2022). These findings indicate that glutamatergic transmission and excitatory pyramidal neuron inputs govern the survival of inhibitory cells in the nascent brain. When migrating GABAergic neurons are devoid of glutamatergic inputs due to the pharmacological blockade of NMDA receptors, inhibition of excitatory neurotransmission has a deleterious influence on immature interneurons and promotes apoptotic death (Roux et al., 2015). Similarly, tetanus toxin injections to ablate excitatory inputs in the somatosensory cortex at postnatal day 0 (P0) caused a notable reduction in the number of somatostatin interneurons in the infragranular cortical layers at P9 (Duan et al., 2020). Intriguingly, the lack of elimination of the excess number of GABAergic neurons during development results in permanent disruptions that only surface in adulthood (Magno et al., 2021). Given the causal observation that the removal of excitatory synaptic inputs during neonatal development at P0 results in interneuron cell death, pyramidal cell activity may act as a key regulator in the emergence of GABAergic circuits.

Studies investigating the role of individual fusion proteins of the SNARE (soluble N-ethylmaleimide fusion protein attachment protein receptor) machinery have confirmed that synaptosomal associated protein 25 kDa (SNAP25)-executed membrane fusion and the regulation of voltage-gated calcium channels are central to synaptic homeostasis and optimal brain functioning. We have previously shown that ablating Snap25 from L5 and L6 projection neurons impacts myelination by reducing the length of the node of Ranvier and the g-ratio in the dorsal column of the spinal cord (Korrell et al., 2019). We have also explored that the presence of regulated vesicle release is fundamental to the maturation of the specialised synapses of L5 corticothalamic projection neurons in the posterior thalamic nucleus (Hayashi et al., 2021) and L5 plays an important part in the regulation of sleep-wake cycles (Krone et al., 2021). It remains to be elucidated; however, how perturbations in pyramidal cell activity impact GABAergic neurons both short-term and long-term. Specifically, how the main output projection neurons of the cortex in layer 5 regulate the distribution of GABAergic neurons in the nascent and the adult brain via activity-dependent mechanisms.

To unveil the role of deep-layer pyramidal neuron activity in orchestrating the spatial and laminar organisation of inhibitory neurons, we selectively manipulated infragranular projection neurons in layer 5 of the cortex. By ablating SNAP25 from selected subsets of glutamatergic L5 projection neurons across the cortical mantle, we abolished Ca^2+-^dependent vesicle release from Rbp4-Cre neurons and therefore, chronically ‘silenced’ L5 projection neurons. The *Rbp4-Cre;Ai14;Snap25^fl/fl^* conditional knockout mice (cKO) allowed us to selectively disrupt Ca^2+^-dependent neurotransmission in infragranular pyramidal neurons while leaving spontaneous and constitutive vesicle release intact (Washbourne et al., 2002). We have previously shown that the presynaptically silenced deep-layer projection neurons follow the same developmental trajectory as that of the control neurons until P21 (Hoerder-Suabedissen et al., 2019). Since no developmental abnormalities have been noticed, the cKO mice thus allow for the genuine investigation of activity-dependent mechanisms on developing GABAergic circuits in a layer-specific fashion. After the first three weeks of development; however, SNAP25 cKO mice exhibit signs of degeneration including impaired axonal integrity and altered synapses, and there is further evidence for altered ultrastructure of neurites, progressive axonal degeneration, and accumulation of inflammation markers in the Snap25-ablated adult brains (Hoerder-Suabedissen et al., 2019). The neurodegenerative processes are only apparent in adult L5-silenced mice; therefore, we could examine how the synaptic activity of L5 impacted the spatial and laminar arrangement of GABAergic neurons *temporarily* (postnatal periods when L5 neurons are intact) and *permanently* (adult stages when L5 neurons start to degenerate). We also distinguished between the *local* (location of the cell bodies) and *global* (subcortical projection sites) effects of chronically silencing cortical L5 neurons on parvalbumin interneurons.

We discovered that the chronic abolition of vesicle release from L5 projection neurons reorganised the laminar distribution of cortical PV neurons in layer 4 of S1 in the adult brain. Interestingly, this effect was exclusive to the adult cortex as the density, distribution, and developmental trajectory of cortical and subcortical PV neurons were unaffected at P14 and P21 suggesting a long-term impact of silencing L5 on the laminar profile of PV cells. Abolishing regulated vesicle release from L5 revealed additional alterations in the laminar positioning of the different subpopulations of PV neurons in L5 (those with and without PNNs), which similarly affected the density of perineuronal nets in the adult motor cortex. Only at P21 did the lack of L5 activity affect the soma features of striatal PV neurons temporarily, as the effect was no longer evident in the adult cortex. We found that the correlation between PV and VVA neurons is influenced by cortical areas and layers and the connection between PV interneurons and perineuronal nets is far more complex than previously thought. Our findings demonstrate that Ca^2+^-dependent synaptic neurotransmission from L5 projection neurons does not impede PV neuron development, but it has a persistent impact on the laminar arrangement of PV neurons, which only manifests in the adult cortex.

## Methods

### Breeding and maintenance of transgenic mice

All experimental procedures were conducted in compliance with the project and personal licenses, as per the rules and regulations of the Animals (Scientific Procedures) Act 1986. The animal work was performed at the Biomedical Services (BMS) of the University of Oxford while adhering to the local regulations related to animal care and welfare. Animals were kept on a 12-hr light/12-h dark cycle and food and water were available *ad libitum*. Transgenic mice with a genetic ablation in the synaptosomal associated protein 25 kDa (SNAP25) gene (B6-Snap25tm3mcw (Snap25-flox))) were used to engineer a cell-type specific conditional knockout strain enabling the selective abolition of Ca^2+^-dependent neurotransmitter release from subsets of glutamatergic projection neurons. The generation and characterisation of the *Rbp4-Cre+;Ai14;Snap25^fl/fl^*mice and the validation of the absence of regulated vesicle release have previously been described (Welch et al., 2000; Marques-Smith et al., 2016; Gustus et al., 2018; Hoerder-Suabedissen et al., 2019; Korrell et al., 2019, Hayashi et al., 2021; Krone et al., 2021).

### Cell-type specific ablation of Snap25 from glutamatergic cortical layer 5 projection neurons

To abolish regulated synaptic vesicle release from subsets of deep-layer glutamatergic neurons, homozygous B6-Snap25tm3mcw (Snap25^fl/fl^) mice were first crossed to B6;129S6-Gt(ROSA)26Sortm14(CAG-tdTomato)Hze/J (Ai14) mice. The homozygous Snap25^fl/fl^;Ai14 mice were then crossed to a bacterial artificial chromosome (BAC) Cre-recombinase driver line Tg(Rbp4-cre)KL100Gsat/Mmucd (Rbp4-Cre; Jackson Laboratories) suitable for the investigation of cortical pyramidal neurons in a layer-specific manner. Cre/+;Snap25^fl/+^;Ai14 female mice were crossed to Snap25^fl/fl^;Ai14 males to generate conditional knockout (*Cre+;Ai14;Snap25^fl/fl^*) and heterozygous knockout (Cre+;Ai14;Snap25^fl/+^) mice. No heterozygous knockout mice were used in the experiments. *Cre-;Ai14;Snap25^fl/+^*and *Cre-;Ai14;Snap25^fl/fl^* were used as ctrl animals. Both male and female mice were used throughout the study. For the developmental studies on the effects of abolishing evoked vesicle release from L5 projection neurons, mice were perfused at postnatal day 14 (P14) and postnatal day 21 (P21) that capture the developmental stages when spontaneous neural activity shapes the emergence of inhibitory cortical circuits. For the adult studies, mice of both sexes were collected at twelve weeks of age where SNAP25 cKO mice already exhibited an accumulation of inflammation markers and signs of axonal degeneration including impaired axonal integrity and altered synapses (Hoerder-Suabedissen et al., 2019; Vadisiute et al., unpublished). For the number of animals used in the study, please see Supplementary Table 1 in the Supplementary materials. For the selected cortical and subcortical regions of interest to distinguish between the local and global effects of L5 on PV neurons, please see Supplementary Table 3.

### Perfusion fixation and vibratome sectioning

Mice aged P14, P21, and 3 months of age were anaesthetised with an overdose of sodium pentobarbital (60 mg/kg, intraperitoneal injection (i.p.)) and were trans-cardially perfused using ice-cold saline (0.9% NaCl) solution followed by 4% paraformaldehyde (PFA, F8775; Sigma-Aldrich) diluted in 0.1 M phosphate-buffered saline (PBS, pH 7.4) (n=3 brains per genotype for P14 experiments (3 ctrl, 3 cKO), n=4 brains per genotype for P21 experiments (4 ctrl, 4 cKO), and n=5 brains per genotype (5 ctrl, 5 cKO) for the 12-week-old adult studies). Brains were dissected and postfixed in 4% PFA for 24 hours at 4^◦^C. Following post-fixation, brains were transferred to 0.1 M PBS containing 0.05% sodium azide (PBSA) (Sodium azide ReagentPlus^®^, S2002-5G, Sigma Aldrich, CAS number: 26628-22-8) and kept at 4^◦^C for long-term storage. Coronal slices of 50 µm thickness were cut from 4.5% agarose-embedded brains using a vibrating microtome (Leica VT1000S; Leica Microsystems, Wetzlar, Germany) and hemisections were collected in 0.1M PBSA in 24-well cell culture plates (CLS3526, Corning^®^ Costar^®^).

### Immunohistochemistry

#### Labelling parvalbumin interneurons and the perineuronal nets

To study how the abolition of Ca^2+^-dependent neurotransmission from selective subsets of L5 pyramidal neurons affects the density of cortical and subcortical GABAergic neurons, double immunohistochemistry (IHC) was performed to label parvalbumin (PV) interneurons and the perineuronal nets (PNNs) using the *Vicia villosa* (VVA) plant lectin. For free-floating PV & VVA double immunostaining, sections were washed three times in 0.1M PBS for 10 minutes. Sections were then permeabilised with a blocking solution containing 2% donkey serum (D9663, Sigma-Aldrich) and 0.2% Triton-X100 (X-100, CAS number: 9002-93-1, Sigma-Aldrich) diluted in VVA buffer for 2 hours at room temperature (RT) prior to overnight incubation with primary antibody at 4^◦^C. For VGLUT1 staining, 5% goat serum and 0.3% Triton-X100 were used. VVA buffer consisted of 0.01 M PBS, 0.15 M NaCl, and 0.1 mM CaCl2 to reach the minimum Ca^2+^ level required for optimal lectin binding. Rabbit anti-PV (Cat# PV27, RRID:AB_2631173, Swant, Marly, Switzerland ) was diluted at 1:5000 for postnatal and 1:500 for adult brains, and the biotinylated VVA lectin (B-1235-2, Vector Laboratories, Burlingame, US) was used at 2 µg/ml concentration and diluted in a blocking solution consisting of 2% donkey serum and 0.2% Triton-X100 diluted in VVA buffer. To ascertain whether Rbp4/tdTomato-expressing axonal processes and VGLUT1-positive synapses colocalize in the striatum, slices were incubated with guinea pig anti-VGLUT1 (1:500, EMD Millipore AB5905, lot: 3258729). After rinsing sections three times in 0.1M PBS, slices were incubated with the following secondary antibodies: goat anti-guinea pig Alexa Fluor^™^ 633 (Invitrogen, A-21105), donkey anti-rabbit Alexa Fluor^™^ 488 (Invitrogen, A-21206, Thermo Fisher, Waltham, MA USA 02451) at 1:500 and Streptavidin-conjugated Cy5 (Invitrogen, SA1011) at 1:200 for 2 hours at RT. Sections were rinsed two times in 0.1M PBS for 10 minutes and counterstained with 4,6-diamidine-2-phenylindole dihydrochloride (DAPI) (1:1000 in 0.1M PBS, overnight). Postnatal coronal sections were coverslipped with 0.1M PBS, while adult sections were coverslipped using Prolong^™^ Gold antifade mountant (P36930, Molecular Probes, Eugene, OR USA 97402). To determine the specificity of the antibody used, control sections were processed following the same steps of immunohistochemistry, but they were incubated only with secondary antibodies. Non-specific staining was not observed in any samples. When performing single immunofluorescence staining for PV, no VVA buffer was used, and sections were permeabilised with a blocking solution containing 2% donkey serum and 0.2% Triton-X100 diluted in 0.1M PBS for 2 hours.

### Image acquisition

Images containing the cortical and subcortical targets of Rbp4-Cre+ L5 projection neurons were acquired with an inverted laser-scanning confocal microscope (Zeiss LSM 710, Germany) equipped with the following laser lines: Diode (405 nm), Argon laser (458, 488, 514 nm), Diode-Pumped Solid-State (DPSS, 561nm), Helium-Neon (633nm). High-resolution confocal images for PV and VVA quantification were acquired using an EC Epiplan-Apochromat 20x (0.80 NA) dry objective at 1x optical zoom, while low-power photomicrographs representing the location of high-power confocal images were obtained with an upright widefield fluorescence microscope equipped with a colour camera for brightfield and fluorescence imaging. (Leica DMR, DFC 280, HC PL FLUOTAR 2.5x / 0.07). For the density and laminar distribution analyses of cortical PV+ neurons, 1x4 tile scans spanning the whole depth of M1 and S1 were acquired using sequential mode scanning. For the density and laminar distribution analyses of cortical VVA+ cells, 1x4 tile scans combined with z-stack imaging were acquired. For subcortical region analyses, 20x confocal image stacks were obtained for PV and VVA quantification and Z-projection images using the maximum intensity function in FiJi (ImageJ) were produced for subsequent cell quantification. For each channel, the optimal laser power, digital gain, offset, and pinhole settings were identified and kept constant throughout image acquisition.

### Image analysis

#### Cell quantification of PV+ neurons

Confocal image files of “.lsm” extensions were processed in FiJi (ImageJ). For the quantification of cortical PV+ interneurons in M1 and S1, a semi-automated custom-designed macro written in FiJi (NIH) was used to assess the density and laminar distribution of PV cells both in the postnatal and adult brains. Due to the varying nature of PV morphology in subcortical brain regions, PV+ cells were quantified manually using the Cell Counter plugin of FiJi (ImageJ) in the globus pallidus, the higher-order thalamic nuclei, and the superior colliculus. The accuracy of the semi-automated cell quantification method was confirmed by manual recounting of cells. The workflow of the custom-designed pipeline is as follows. In brief, multichannel confocal images were first duplicated and split into individual channels using the FiJi Color -> Split Channels function. Pre-processing of images was then conducted using the rolling ball algorithm to correct uneven backgrounds and a linear Gaussian blur filter with a sigma value of 1 was applied to reduce noise. After filtering and noise correction, segmentation of foreground (objects of interest) from background objects was performed using the Otsu thresholding method. Following segmentation, a Watershed binary process was applied to separate touching or merged objects. Finally, the Analyse particle > Set measurements function was used to set the size and circularity of objects (size=40-Infinity, circularity=0.10-1.00). Objects falling outside the preset range were excluded from the cell measurements. Double-positive PV+ VVA+ cells were identified based on the colocalization of PV with the VVA channel and were manually counted.

### Cell quantification of VVA+ cells

For the density and laminar distribution analysis of VVA+ cells in the adult brain, manual cell counting was selected over semi-automated quantification as the segmentation of the lattice-like morphology of PNNs proved to be unreliable. Manual quantification of VVA+ cells was performed using the Cell Counter plugin of ImageJ. VVA+ cells that exhibited a typical web-or lattice-like morphology and co-localised with the nuclear counterstain DAPI signal were counted as positives. Immunohistochemical staining of PNNs in the developing brains yielded varied results and therefore, the VVA+ cells were only quantified in the adult brain at 3 months of age. In the postnatal brains, the visualization of PNNs was either insufficient due to low staining intensity or the staining produced diffuse non-specific labelling in subcortical structures. The typical web or lattice-like structure of PNNs was not detectable in all postnatal brains. For the purpose of quantifying the density of VVA+ cells in nascent brains, P14 and P21 brains were deemed inappropriate. Considering the condensation of proteoglycans into PPNs is not yet complete at such early developmental ages, the detection of PNNs and the quantification of VVA+ cells might not be feasible with plant lectins during early brain development.

### Morphometric analysis of PV interneurons

Point-scanning confocal z-stack images of the caudoputamen were obtained using a 20x objective (Epiplan-Apochromat 20x (0.80 NA) dry) and maximum-intensity projected in Fiji to construct 2D images for morphometric analysis of PV neurons. The analysis pipeline was identical to that of the density and laminar distribution analyses, and the Analyse > Set measurements function was used to select the parameters to be investigated, i.e., *soma area, circularity, perimeter, feret, minferet, feret angle, roundness, solidity*. Following the automated quantification of PV cells, the result of density counts, and morphometric measurements were stored in “.csv” files and exported to Prism 10 (GraphPad) for plotting and statistical analysis. Scatter plots of morphometric analyses of PV+ cells represent individual cell measurements. For the number of cells measured for morphometric analyses and the number of cells examined at different time points, please see Supplementary Table 2.

### Statistics

Cell density and laminar distribution analyses of PV+ and VVA+ were performed on data obtained from n=3 brains per genotype for P14 experiments (3 ctrl, 3 cKO), n=4 brains per genotype for P21 experiments (4 ctrl, 4 cKO), and n=5 brains per genotype (5 ctrl, 5 cKO) for the 12-week-old adult studies. Box plot values are represented as mean ± SEM. When comparing two groups (*morphometric analysis of PV+*), a two-tailed unpaired t-test with Welch’s correction was applied. When comparing more than two groups (density, distribution, and trajectory of PV+ and VVA+ cells), a two-way ANOVA with Šídák’s multiple comparisons test was used. Scatter plots of morphometric analyses of PV+ cells represent individual cell measurements. Heatmaps present the Pearson correlation between PV and VVA in different cortical regions and different cortical layers (PV, n=5 per genotype; VVA, n=4 per genotype). Statistical tests were computed in Prism 10.2.1 (GraphPad Software, San Diego, CA, USA).

## Results

### The Rbp4-Cre;Snap25^fl/fl^ mice genetically label two types of cells - cortical layer 5 projection neurons and hippocampal granule cells

To elucidate the role of glutamatergic layer 5 cortical projection neurons in the development and spatial organisation of GABAergic PV neurons, we conditionally deleted *SNAP25* from Rbp4-Cre+ neurons in L5 of the cortex and thus selectively abolished calcium-dependent vesicle release from specific subsets of L5 neurons. To distinguish between the *local* vs *global* effects of L5 on the density and distribution of PV interneurons, we selected our regions of interest (ROI) based on the location of the cell bodies (*local effec*t) and the long-range axonal projections of the silenced L5 Snap25 cKO neurons (*global effect*). The primary motor (M1) and the primary somatosensory cortices are densely populated with Rbp4-Cre+ cortical projection neurons (Figure 1 A2, A6, A10) and PV interneurons (Figure 1 A3, A7, A11); therefore, we selected M1 and S1 as our cortical ROIs. In the *Rbp4-Cre+;Ai14;Snap25^fl/fl^* mice, Cre expression is mostly detected in the pyramidal tract (PT) and intratelencephalic (IT) subgroups of L5 projection neurons throughout the neocortex. However, Cre-expressing tdTomato+ neurons were also observed in the dentate gyrus (Figure 1 B2, B4, B6, B8, B10, B12) indicating that Ca^2+^-dependent vesicle release was ablated in the granule cells in addition to the Rbp4-Cre+ L5 neurons. The Snap25 cKO mice can thus be used not only for studying the effects of chronically silencing L5 projection neurons but also for the effects of chronically silencing the mossy fibre pathway on PV interneurons (Figure 1 B3, B7, B11).

**Figure 1.**
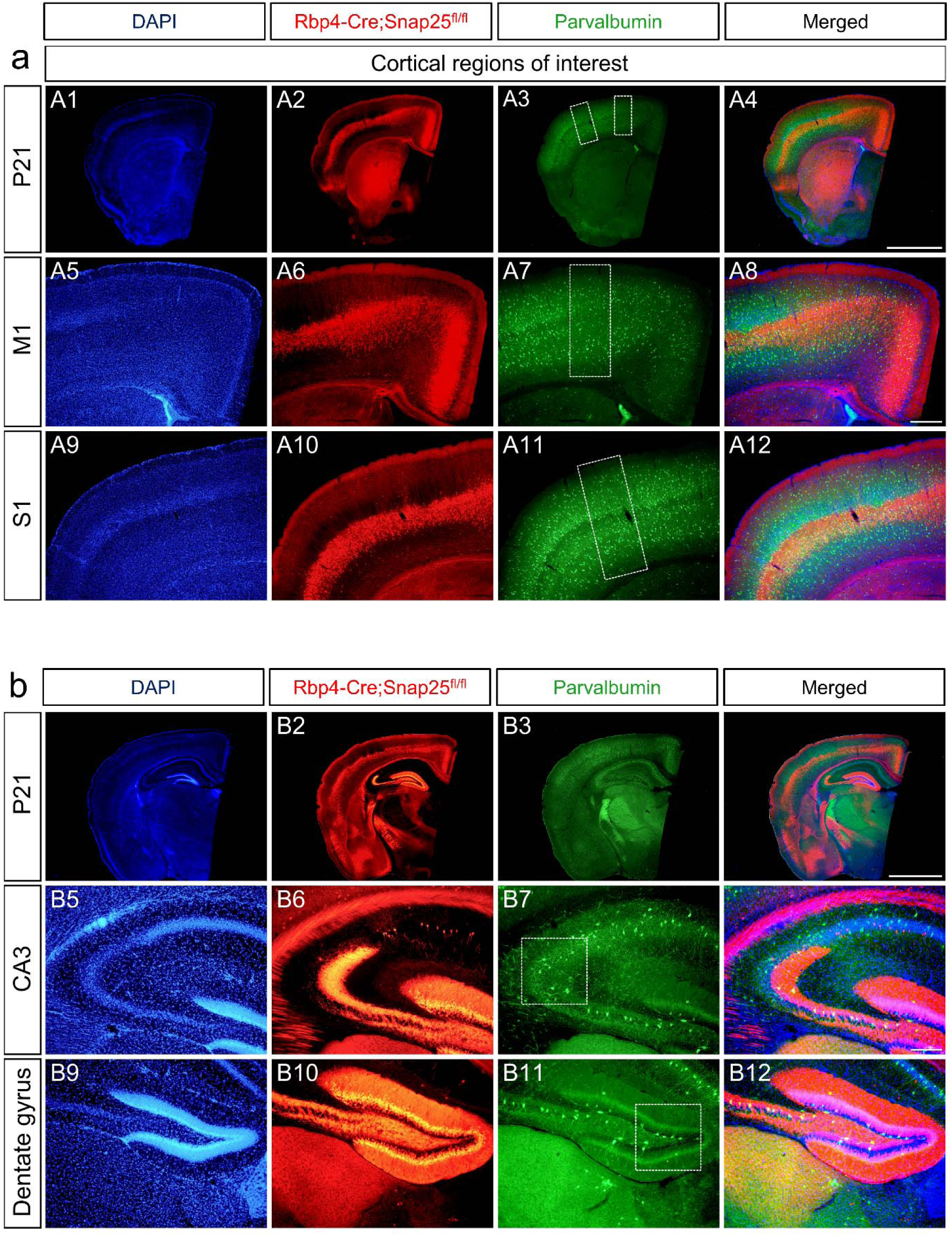
Distribution of Rbp4-Cre-expressing neurons in layer 5 of the cortex and the dentate gyrus of the hippocampus in the Snap25 cKO mice. Low-magnification epifluorescence photomicrographs of Snap25 cKO brains at P21 (A1-A4, B1-B4) counterstained with DAPI (blue) and immunostained for parvalbumin neurons (green) (A3, B3). Epifluorescence images illustrating the distribution of silenced Rbp4-Cre+ projection neurons in the primary motor and somatosensory cortices (A6, A10). Note that the Rbp4-Cre driver line labels mixed subpopulations of intratelencephalic and extratelencephalic projection neurons in L5 of the cerebral cortex (A2, B2). Merging *Rbp4-Cre;Ai14; Snap25^fl/fl^* and PV epifluorescence micrographs reveals the presence of PV-immunoreactive neurons in the vicinity of the cell bodies of L5 projections neurons in the primary motor and somatosensory cortices (A8, A12). Cortical regions of interest (ROIs) selected to study the local effects of abolishing evoked vesicle release from L5 pyramidal neurons on PV interneurons are marked by white boxes (A7, A11). Nuclear counterstains of cortical and hippocampal ROIs (A5, A9, B5, B9). Epifluorescence images depicting the distribution of Cre expression in the developing Snap25 cKO brains in L5 across the cortical mantle and the hippocampus (B2, B4). Note the robust Cre expression in the granule cells of the dentate gyrus indicating the chronic silencing of the mossy fibre pathway (B6, B10). White boxes denote hippocampal ROIs, i.e., CA3 and dentate gyrus (B7, B11). Fluorescence micrographs of Cre-expressing hippocampal granule cells superimposed on PV+ interneurons in the Snap25 cKO mice (B4, B8, B12). Scale bars: 1000 µm (A4, B4), 200 µm (A8), 100 µm (B8).

### Chronically ‘silenced’ Rbp4-Cre+ glutamatergic projection neurons in layer 5 exhibit similar corticothalamic and corticofugal projections as Rbp4-Cre+ neurons

To investigate if the chronic removal of Ca^2+-^dependent neurotransmission from L5 has a global effect on PV neurons, we selected those output regions of Rbp4-Cre+ tdTomato+ L5 neurons where PV-immunoreactive cells were present (Figure 2 A3, A7, A11, A15, A19, A23). Rbp4-Cre;Snap25^fl/f^ L5 neurons project to the contralateral caudoputamen (CPu) (Figure 2 A2, A4), the globus pallidus external segment (GPe) (Figure 2 A6, A8), the higher-order thalamic nuclei including the lateral posterior nucleus (LP) (Figure 2 A10, A12) and the mediodorsal nucleus of the thalamus (MD (Figure 2 A14, A16). Collaterals of the silenced Rbp4-Cre L5 axons also target midbrain areas such as the pontine nuclei (Figure 2 A22, A24) and the superior colliculus (SC) (Figure 2 A18, A20), and they extend to the spinal cord through the pyramidal tract. Of the output regions of L5 Rbp4-Cre;Snap25^fl/fl^ neurons, the following innervation sites were selected as our subcortical ROIs: CPu, LP, MD, GPi, and SC (Supplementary Table 3). These subcortical regions are of great importance as the silenced pyramidal cells of layer 5 could exert a considerable effect on the number and distribution of PV+ cells through their corticofugal projections.

**Figure 2.**
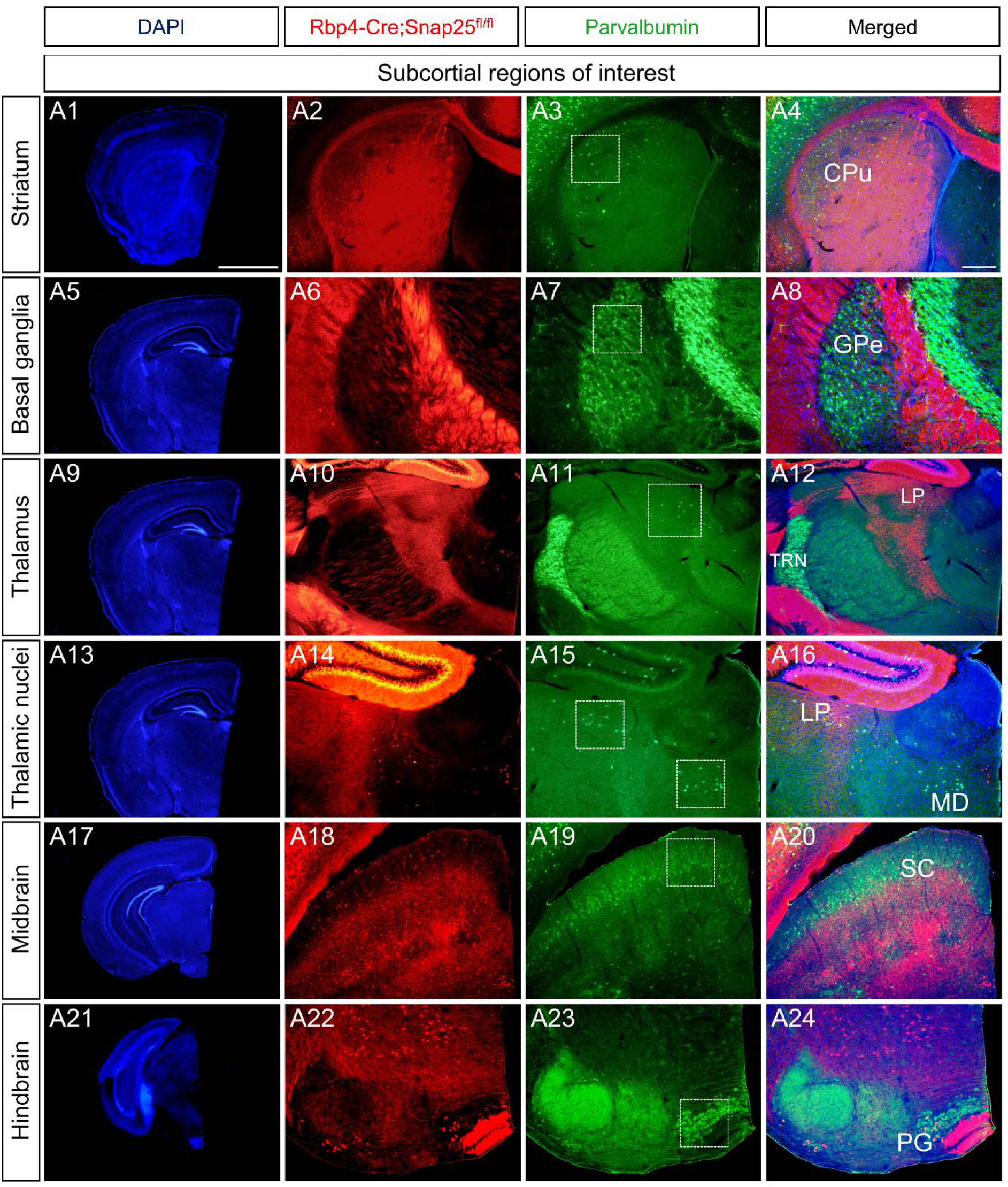
Long-range axonal projections of the ‘silenced’ Rbp4-Cre;Snap25^fl/fl^ projection neurons in layer 5 of the cerebral cortex. Epifluorescence micrographs of coronal hemisections of P21 Snap25 cKO mice obtained at different rostrocaudal levels and counterstained with DAPI (A1, A5, A9, A13, A17, A21). Long-range corticothalamic and subcortical axonal projections of *Rbp4-Cre;Ai14; Snap25^fl/fl^* neurons innervate the basal ganglia (A2, A6), the higher-order thalamic nuclei (A10, A14), and the midbrain superior colliculus (A18). Collaterals of the descending corticofugal fibres of Rbp4-Cre+ L5 projection neurons target the brainstem motor regions including the pontine gray (A22) before reaching the spinal cord. White boxes define subcortical regions of interest (ROIs): the caudoputamen (A2-A4), the external segment of the globus pallidus (A6-A8), the lateral posterior nucleus of the thalamus (A10-A12, A15-16), the mediodorsal thalamic nucleus (A15-A16), the superior colliculus (A18-A20), and the pontine gray (A22-A24). Specific subcortical projection sites of Rbp4-Cre+ L5 neurons were carefully selected to explore the global effects of abolishing Ca^2+-^dependent neurotransmission from L5 projection neurons. Note the distribution of subcortical PV neurons overlaps with the output regions of silenced L5 cortical projection neurons (A4, A8, A12, A16, A20, A24). Scale bars: 1000 µm (A1), 200 µm (A4). HO nuclei: higher-order nuclei.

### Chronic abolition of evoked vesicle release from layer 5 projection neurons does not interfere with the development of PV neurons in the second and third weeks of postnatal development

To ascertain whether regulated synaptic vesicle release from L5 is required for the development of PV interneurons, we assessed the local effects of silencing Rbp4-Cre+ neurons on the density of cortical PV neurons at postnatal day 14 (P14) and postnatal day 21 (P21). No changes were detected in the density of PV-immunoreactive cells in M1 at P14 and P21 (Figure 3 A1-A1’, B1-B1’, E1, E1’, E3, E3’, f; two-way ANOVA with Šídák’s multiple comparisons test, p= 0.3679 (P14), p= 0.6323 (P21)). At both P14 and P21, the density of cortical PV neurons in S1 remained unaffected indicating the abolition of evoked neurotransmitter release from L5 pyramidal neurons does not influence the density of cortical PV+ cells (Figure 3 C1-C1’, D1-D1’, E2, E2’, E4, E4’, g; two-way ANOVA with Šídák’s multiple comparisons test, p=0.5806 (P14), p=0.8327 (P21)). After confirming that there were no changes in the density of PV neurons in M1 and S1, we investigated the developmental trajectory of cortical PV interneurons during P14 and P21. Neither M1 nor S1 showed a change in PV cell density between P14 and P21 in the ctrl brains suggesting that there is no significant decrease in the number of PV interneurons following developmentally programmed cell death (Figure 3h, two-way ANOVA with Šídák’s multiple comparisons test, p= 0.7467 (M1 ctrl), p= 0.8398 (S1 ctrl)). The developmental trajectory of cortical PV+ interneurons in the mutant brains was identical to the trajectory observed in the ctrl brains (Figure 3i, two-way ANOVA with Šídák’s multiple comparisons test, p= 0.9868 (M1 cKO), p= 0.9671 (S1 cKO)). These results suggest that the development of cortical PV interneurons is not influenced by the chronic abolition of neurotransmitter release from L5 projection neurons at P14 and P21.

**Figure 3.**
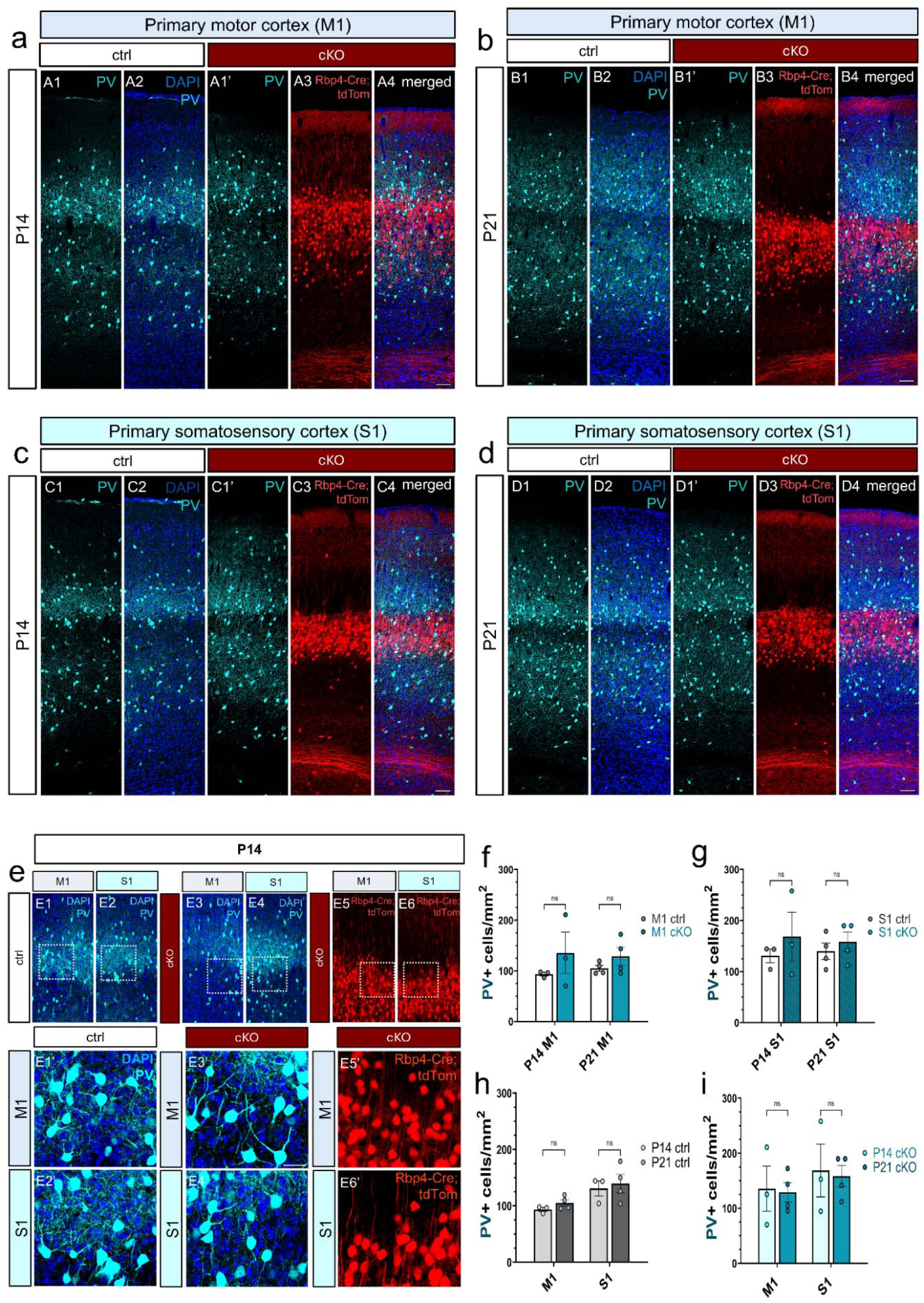
Perturbing the activity of Rbp4-Cre+ projections neurons in cortical layer 5 has no local effect on cortical PV interneurons. 2D tile scans of point-scanning confocal microscope images of the primary motor cortex during the second and third postnatal weeks of development. PV-immunoreactive neurons (A1-A1’, B1-B1’) and Rbp4-Cre;Snap25^fl/fl^ projection neurons in the L5-silenced Snap25 cKO brains in M1 at P14 (A3-A4) and P21 (B3-B4). Laser-scanning confocal tile-scan images of the primary somatosensory cortex depicting PV-positive neurons and Rbp4-Cre;Snap25^fl/fl^ projection neurons (C3-C4, D3-D4) in S1 in the ctrl and Snap25 cKO brains at P14 (C1-C1’) and P21 (D1-D1’). High-magnification confocal images of PV interneurons in M1 and S1 in the ctrl (E1, E2, E1’, E2’) and L5-silenced brains at P14 (E3, E4, E3’,E4’). Snap25-abolished cortical projection neurons in layer 5 of M1 and S1 at P14 (E5, E6, E5’, E6’). The regions of high-magnification images are delineated by dotted white boxes. Quantification of PV cells in M1 in the second and third postnatal weeks revealed no significant differences in the density of PV neurons between the ctrl and the chronically silenced L5 brains (f). No changes were detected in the density of PV neurons in S1 at P14 and P21 either (g). The developmental trajectory of cortical PV interneurons in the ctrl brains revealed no significant changes in their density between P14 and P21 in the two cortical regions investigated (h). Abolition of regulated vesicle release from L5 pyramidal neurons did not alter the developmental trajectory of PV neurons (i). Scale bars: 100 µm (all panels). Data is represented as mean ± SEM. M1: primary motor cortex; S1: primary somatosensory cortex.

### Chronic abolition of evoked vesicle release from layer 5 projection neurons leaves subcortical PV interneurons intact during the early stages of development

Next, we assessed the global effect of the chronic abolition of regulated vesicle release from L5 on subcortical PV neurons. At P14, none of the subcortical regions displayed a difference in the density of PV-immunoreactive cells between the ctrl and the Snap25 cKO brains. (Figure 4a A1-10, 4c) (LP: 50.4 ± 5.9 (ctrl), 64.4 ± 28.5 (cKO), p=0.999; MD: 59.6 ± 4.6 (ctrl), 48.0 ± 5.2 (cKO), p= >0.9999; CPu: 84.4 ± 6.6 (ctrl), 69.7 ± 3.4 (cKO), p=0.9998; GPe: 381.1 ± 39.4 (ctrl), 420.3 ± 71.8 (cKO), p= 0.9813; SC: 246.6 ± 40.3 (ctrl), 355.3 ± 106.8 (cKO), p= 0.4308; all two-way ANOVA with Šídák’s multiple comparisons test). Similarly, no alterations were found in the density of subcortical PV interneurons at P21 implicating the chronic cessation of layer 5 activity since birth does not have a global influence on the development of PV neurons (Figure 4b B1-B10, 4d) (LP: 50.2 ± 14.0 (ctrl), 63.6 ± 10.0 (cKO), p= >0.9999; MD: 62.8 ± 1.6 (ctrl), 64.3 ± 10.2 (cKO), p= >0.9999; CPu: 86.3 ± 5.1, (ctrl), 85.3 ± 4.9 (cKO), p= >0.9999; GPe: 397.0 ± 50.8 (ctrl), 485.0 ± 117.4 (cKO), p= 0.7598; SC: 438.7 ± 94.8 (ctrl), 415.7 ± 45.9 (cKO), p= 0.9992; all two-way ANOVA with Šídák’s multiple comparisons test). Considering that the maturation of subcortical PV+ neurons may follow a different trajectory than that of the cortical PV+ cells, we next established the developmental profile of subcortical PV neurons. Of the output regions of L5 projection neurons, there was a significant increase in the density of PV+ cells in the superior colliculus between P14 and P21 in the ctrl brains (Figure 4e) (P14: 246.6 ± 40.3 (ctrl), P21: 438.7 ± 94.8 (ctrl), p= 0.0217, two-way ANOVA with Šídák’s multiple comparisons test). No changes were observed in the higher-order thalamic nuclei and the basal ganglia (Figure 4e) (P14 LP: 50.4 ± 5.9 (ctrl), P21 LP: 50.2 ± 14.0 (ctrl), p= >0.9999; P14 MD: 59.6 ± 4.6 (ctrl), P21 MD: 62.8 ± 1.6 (ctrl), p= >0.9999; P14 CPu: 84.4 ± 6.6 (ctrl), P21 CPu: 87.0 ± 5.4 (ctrl), p= >0.9999; P14 GPe: 381.1 ± 39.4 (ctrl), P21 GPe: 397.0 ± 50.8 (ctrl), p= 0.9997, all two-way ANOVA with Šídák’s multiple comparisons test). As opposed to the significant rise noted in the ctrl brains, PV cell density did not follow the same trend in the Snap25 cKO mice between P14 and P21 (Figure 4f) indicating a different developmental trajectory of PV neurons in the superior colliculus after L5 was silenced (P14: 355.3 ± 106.8 (cKO), P21: 415.7 ± 45.9 (cKO), p= 0.9605). There were no alterations in PV cell density in the thalamic nuclei and the basal ganglia either in the layer 5-silenced brains (Figure 4f) (P14 LP: 64.4 ± 28.5 (cKO), P21 LP: 63.6 ± 10.0 (cKO), p= >0.9999; P14 MD: 48.0 ± 5.2 (cKO), P21 MD: 64.3 ± 10.2 (cKO), p= >0.9999; P14 CPu: 69.7 ± 3.4 (cKO), P21 CPu: 85.3 ± 4.9 (cKO), p= >0.9999; P14 GPe: 420.3 ± 71.8 (cKO), P21 GPe: 485.0 ± 117.3 (cKO), p= 0.9474, all two-way ANOVA with Šídák’s multiple comparisons test).

**Figure 4.**
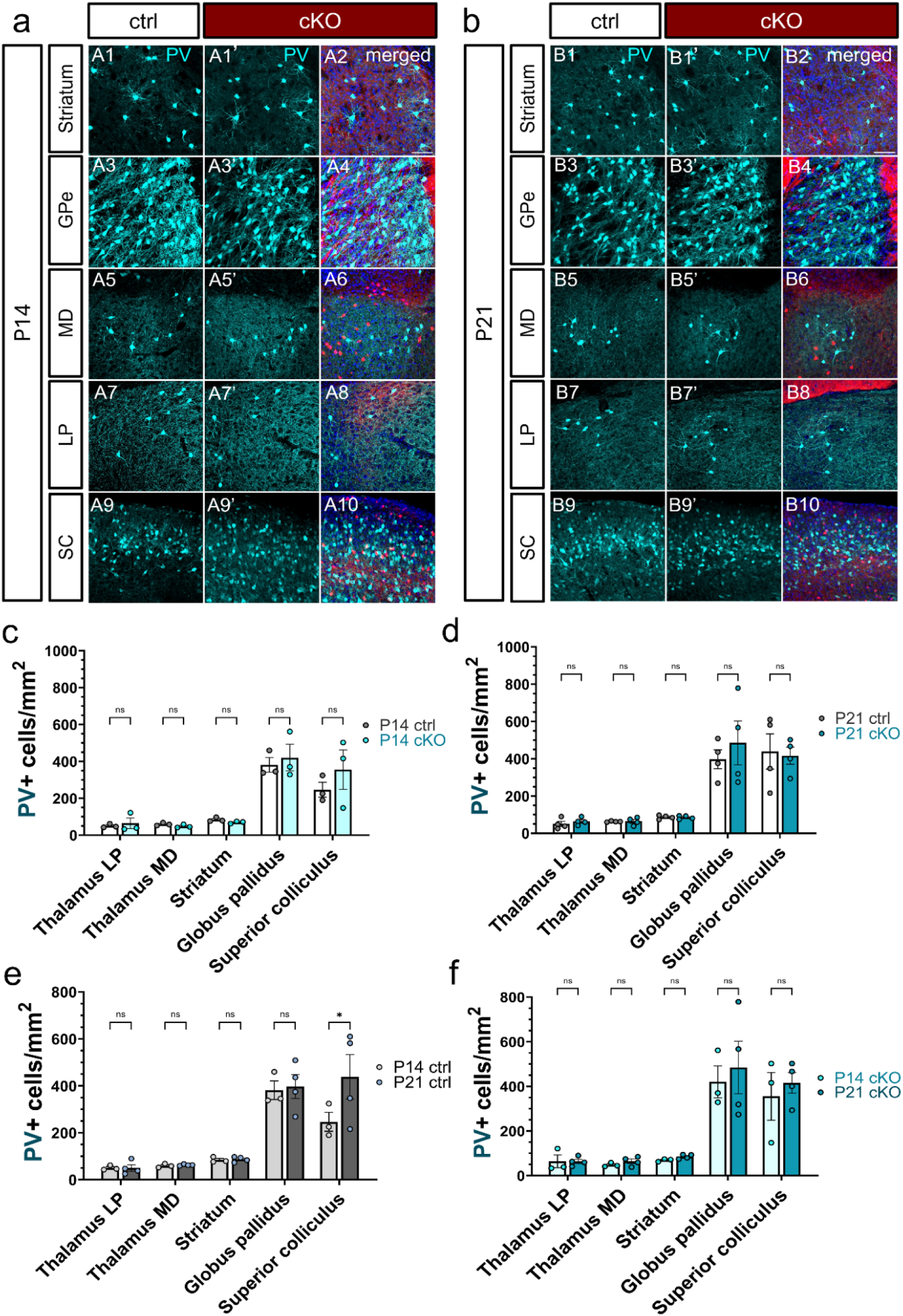
Perturbing the activity of Rbp4-Cre+ projections neurons in cortical layer 5 has no global impact on subcortical PV interneurons. Maximum intensity projection of confocal z-stack images representing the output regions of cortical Rbp4-Cre+ L5 projection neurons at various rostrocaudal levels in the Snap25 ctrl and cKO brains in the second and third postnatal week. Parvalbumin-immunoreactive neurons (cyan) shown in the caudoputamen (A1-A2) and the external segment of the globus pallidus (A3-A4) in the L5-silenced ctrl and cKO brains at P14 (A1-A4) and P21 (B3-B4). PV+ neurons in the higher-order lateral posterior and mediodorsal nuclei of the thalamus at P14 (A5-A8) and P21 (B5-B8) which are selectively innervated by cortical L5 and L6b glutamatergic projection neurons. Maximum-intensity projected z-stack image showing PV-immunoreactive neurons in the superior colliculus at P14 (A9-10) and P21 (B9-B10). Quantification of PV+ cell density in the subcortical projection regions of *Rbp4-Cre;Ai14;Snap25^fl/fl^* cortical neurons at P14 and P21 revealed no significant differences in the density of PV+ neurons between the Snap25 ctrl and cKO brains (c, d). The developmental trajectory of subcortical PV neurons in the output regions of L5 neurons assessed between P14 and P21 in the ctrl and cKO brains (e, f). The ctrl brains showed a significant increase in the density of PV+ neurons in the superior colliculus (e) between P14 and P21, whereas no such change was observed in the L5-silenced mice (f). Scale bars: 100 µm. All data is represented as mean ± SEM values. *p < 0.05, p = 0.0217. 2-way ANOVA with Šídák’s multiple comparisons test.

### The chronic absence of synaptic vesicle release from Rbp4-Cre+ layer 5 pyramidal neurons causes alterations in the morphology of striatal PV+ neurons at P21

After establishing that the chronic cessation of Ca^2+^-dependent neurotransmission from Rbp4-Cre+ L5 projections neurons does not have a global influence on the density of subcortical PV+ neurons in its projection regions, we next investigated whether the disruption of L5 affects the morphology of PV cells in the striatum. This output region of L5 is not only innervated by the axonal projections of Rbp4-Cre+ neurons (Figure 5c C3, C3’, 9e E3, E3’) but also receives dense synaptic innervation as shown by the presence of Vglut1+ tdTomato+ punctae in the dorsal striatum (Figure 5a, b). When assessing the effect of chronically silencing L5 projection neurons on the morphology of striatal PV+ neurons, no differences were detected between the ctrl and the cKO brains at P14 in any of the measured parameters of striatal PV+ neurons (Figure 5c, d) (soma area: 172.6 ± 5.5 (ctrl), 174.7 ± 6.4 (cKO), p= 0.8085; perimeter: 58.6 ± 1.6 (ctrl), 60.5 ± 2.3 (cKO), p= 0.5071; circularity: 0.7 ± 0.0 (ctrl), 0.7 ± 0.0 (cKO), p= 0.7071; feret: 20.4 ± 0.5 (ctrl), 20.7 ± 0.6 (cKO), p= 0.6807; feret angle: 108.1 ± 3.8 (ctrl), 112.8 ± 4.2 (cKO), p= 0.4092, minferet: 13.4 ± 0.3 (ctrl), 13.7 ± 0.4 (cKO), p= 0.4316; roundness: 0.7 ± 0.0 (ctrl), 0.7 ± 0.0 (cKO), p= 0.2717; solidity: 0.9 ± 0.0 (ctrl), 0.9 ± 0.0 (cKO), p= 0.2479, all unpaired t test with Welch’s correction). However, at P21, we noticed a marked increase in the feret angle and a significant decrease in the soma area, minferet, and roundness of striatal PV+ neurons in the L5-silenced brains (Figure 5e, f) (soma area: 161.7 ± 4.7 (ctrl), 147.1 ± 4.0 (cKO), p= 0.0192; perimeter: 57.9 ± 1.6 (ctrl), 55.3 ± 1.4 (cKO), p= 0.2311; circularity: 0.6 ± 0.0 (ctrl), 0.6 ± 0.0 (cKO), p= 0.9626; feret: 20.1 ± 0.5 (ctrl), 19.6 ± 0.4 (cKO), p= 0.4288; feret angle: 98.6 ± 3.9 (ctrl), 109.0 ± 3.3 (cKO), p= 0.0436; minferet: 13.1 ± 0.3 (ctrl), 12.2 ± 0.2 (cKO), p= 0.0158; roundness: 0.7 ± 0.0 (ctrl), 0.7 ± 0.0 (cKO), p= 0.0035; solidity: 0.9 ± 0.0 (ctrl), 0.9 ± 0.0 (cKO), p= 0.7592, all unpaired t test with Welch’s correction). Although the chronic abolition of vesicle release from L5 did not alter the density and the developmental trajectory of striatal PV+ neurons at P21, it did modify their morphology. Striatal PV+ interneurons appeared to be more elongated, with less rounded and smaller somata in the layer 5-silenced mice.

**Figure 5.**
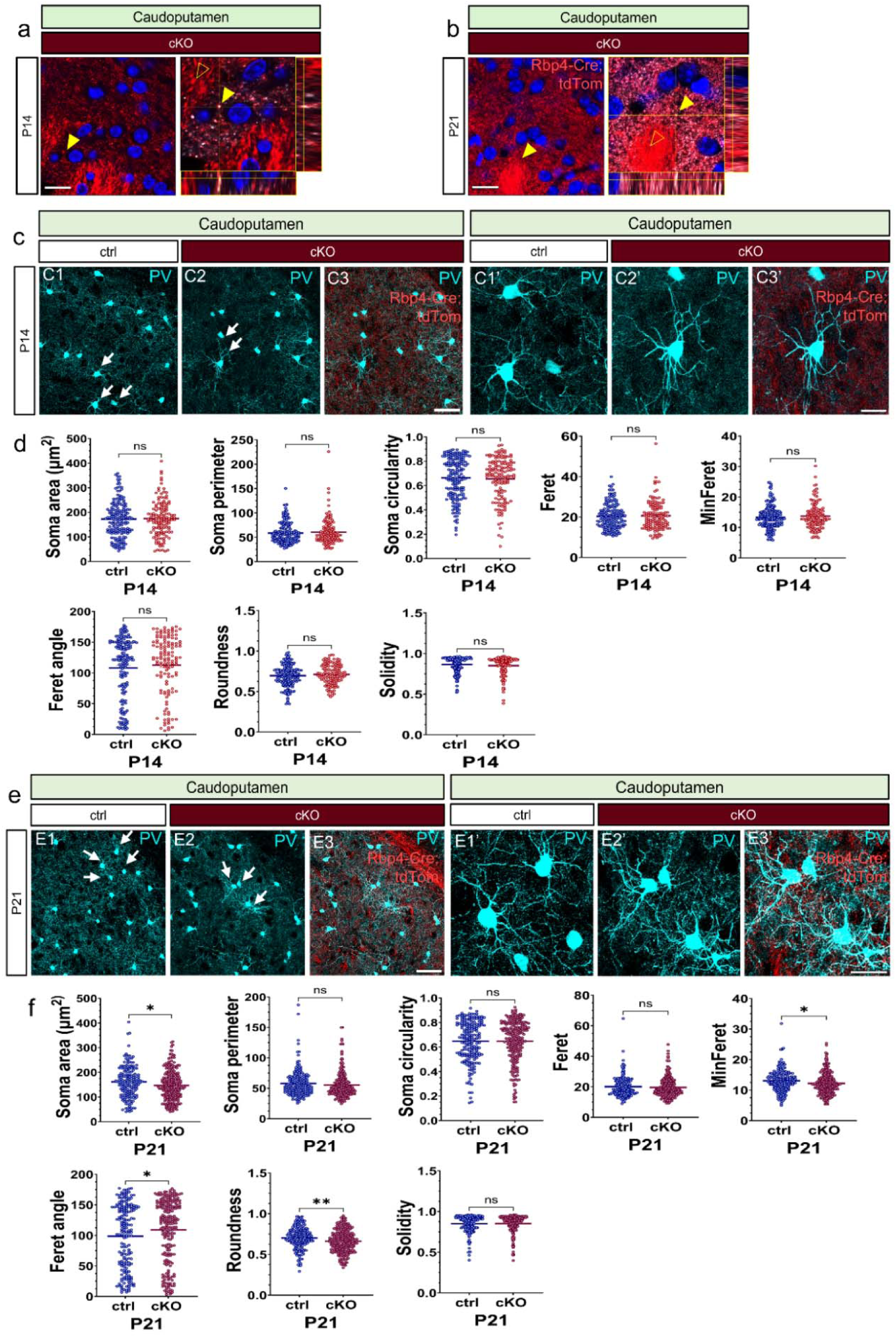
Abolition of Ca^2+-^dependent neurotransmission from layer 5 pyramidal neurons temporarily alters the morphology of striatal PV interneurons in the juvenile brain. Glutamatergic synapses are formed in the dorsal striatum by Rbp4-tdTomato+ neurons (a, b). Confocal point-scanning images of P14 and P21 *Rbp4-Cre;Ai14;Snap25^fl/fl^* brains displaying puncta of VGluT1+ and tdTomato+ in the dorsal striatum in addition to orthogonal views through image stacks. Fluorescence colocalization is evident at both ages (yellow arrowheads), with more punctae noted at P21 (white arrowheads). The tdTom+ fibre bundles (empty yellow arrowheads) are negative for VGluT1 staining (a, b). Maximum-intensity projections of confocal z-stack images of PV-immunoreactive neurons in the caudoputamen at P14 and P21 in the ctrl (C1, C1’, E1, E1’) and the L5-silenced brains (C2, C2’, E2, E2’). Merged maximum-intensity projected confocal images demonstrating the presence of tdTomato+ fibres in the caudoputamen in the vicinity of PV+ neurons (C3, C3’, E3, E3’). There were no significant alterations in the assessed soma features of PV+ neurons as shown by the morphometric analyses of PV+ interneurons at P14 (d). Significant reductions in the soma area, the minferet, and the soma roundness of striatal PV+ neurons were detected at P21 in the Snap25 cKO mice (f) (soma area, p= 0.0192; minferet, p= 0.0158; roundness, p= 0.0035, ctrl, n=180; cKO, n=236 cells). The feret angle of striatal PV+ cells was significantly increased in the L5-silenced mice at P21 (p= 0.0436). Unpaired t-test with Welch’s correction. Scale bars: 50 µm (C3, E3, E3’), 20 µm (a, b, C3’).

### Striatal PV interneurons in the layer 5-silenced brains display altered developmental trajectory of soma features between postnatal stages of development

Having shown the chronic cessation of regulated vesicle release from L5 projection neurons causes alterations in the soma size, feret diameter, and roundness of striatal PV+ neurons at P21, we next assessed the course of striatal PV+ morphology between P14 and P21 in the ctrl brains. None of the examined characteristics of the morphology of PV neurons in the striatum showed significant alterations between P14 and P21 (Figure 6a, b). In the layer 5-silenced brains; however, we revealed a significant reduction in the striatal PV+ neurons’ soma area, minferet, and roundness between 14 and P21 (Figure 6c, d) (soma area: 175 ± 6.4 (P14, cKO), 147 ± 4.0 (P21, cKO), p= 0.0003; perimeter: 60.5 ± 2.3 (P14, cKO), 55.3 ± 1.4 (P21, cKO), p= 0.0558; circularity: 0.7 ± 0.0 (P14, cKO), 0.6 ± 0.0 (P21, cKO), p= 0.6464; feret: 20.7 ± 0.6 (P14, cKO), 19.6 ± 0.4 (P21, cKO), p= 0.1402; feret angle: 113 ± 4.2 (P14, cKO), 109 ± 3.3 (P21, cKO), p= 0.4776; minferet: 13.7 ± 0.4 (P14, cKO), 12.2 ± 0.2 (P21, cKO), p= 0.1402; roundness: 0.7 ± 0.0 (P14, cKO), 0.7 ± 0.0 (P21, cKO), p= 0.0003; solidity: 0.9 ± 0.0 (P14, cKO), 0.9 ± 0.0 (P21, cKO), p= 0.9262, all unpaired t test with Welch’s correction). The somata of striatal PV+ cells in the Snap25 cKO mice became more compressed, smaller, and less rounded due to the considerable decrease in their soma area, roundness, and minferet values. Given that no such morphological alterations were noted in the trajectory of PV+ neurons in the ctrl striatum between P14 and P21, the altered morphology of striatal PV+ neurons in the cKO brains seemed to be caused by the chronic abolition of synaptic transmission from L5.

**Figure 6.**
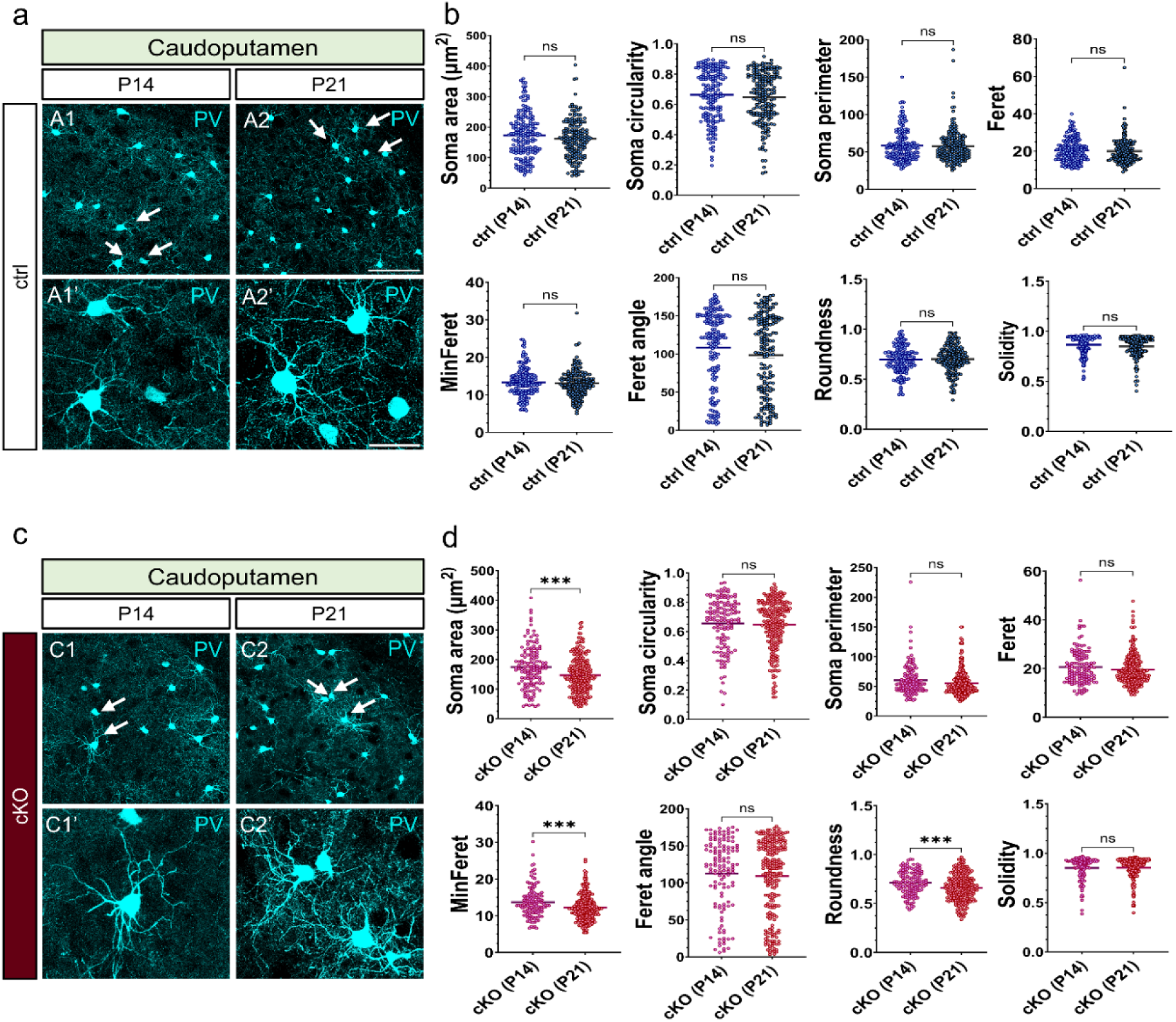
Striatal PV interneurons in the layer 5-silenced brains display altered developmental trajectory of soma features between early stages of development. Maximum-intensity projected confocal z-stack images of PV-immunoreactive neurons in the caudoputamen in the ctrl mice at P14 (A1, A1’) and P21 (A2, A2’). The trajectory of PV neuron morphology in the caudoputamen of ctrl mice did not exhibit any significant alterations in the measured soma characteristics of PV+ neurons between P14 and P21 (b). Maximum-intensity projected confocal z-stack images of PV-immunoreactive neurons in the caudoputamen in the Snap25 cKO mice at P14 (C1, C1’) and P21 (C2, C2’). In the L5-silenced animals, there was a considerable reduction in the soma area, minferet, and roundness of striatal PV+ interneurons between P14 and P21 (d) (soma area, p= 0.0003, minferet, p= 0.0005, roudness, p= 0.0003. Unpaired t test with Welch’s correction. P14, n=136, P21, n=236). Arrows denote PV+ neurons shown in high-magnification images. Scale bars: 100 µm (A2), 20 µm (A2’).

### Chronic impairments in synaptic vesicle release from layer 5 projection neurons have no long-term, permanent impact on the morphology of striatal interneurons in the adult brain

To further investigate if the perceived changes in the morphology of PV interneurons at P21 persist into adulthood, we conducted the same morphometric analyses on striatal PV neurons in the adult brain. Interestingly, none of the observed changes in PV morphology were present at three months of age in the L5-silenced brains (Figure 7a, b) (soma area: 121 ± 3.8 (ctrl), 129 ± 4.7 (cKO), p= 0.2301; perimeter: 56.8 ± 1.9 (ctrl), 56.6 ± 2.1 (cKO), p= 0.9398; circularity: 0.6 ± 0.0 (ctrl), 0.6 ± 0.0 (cKO), p= 0.1721; feret: 19.2 ± 0.5 (ctrl), 19.4 ± 0.6 (cKO), p= 0.7569; feret angle: 92.0 ± 4.2 (ctrl), 96.7 ± 4.1 (cKO), p= 0.4261; minferet: 11.6 ± 0.3 (ctrl), 11.9 ± 0.3 (cKO), p= 0.3774; roundness: 0.7 ± 0.0 (ctrl), 0.7 ± 0.0 (cKO), p= 0.4804; solidity: 0.8 ± 0.0 (ctrl), 0.8 ± 0.0 (cKO), p= 0.3412, all unpaired t test with Welch’s correction). After establishing that the chronic cessation of regulated vesicle release from L5 causes temporary changes in the morphology of striatal PV neurons, we assessed how the trajectory of PV morphology changed from development to adulthood. The striatal PV neurons in the ctrl brains exhibited distinct alterations in soma area, circularity, minferet, roundness, and solidity. By three months of age, all these characteristics had significantly decreased (Figure 7c, d) (soma area: 162 ± 4.7 (P21, ctrl), 121 ± 3.8 (12 wks, ctrl), p= <0.0001; perimeter: 57.9 ± 1.6 (P21, ctrl), 56.8 ± 1.9 (12 wks, ctrl), p= 0.6702; circularity: 0.6 ± 0.0 (P21, ctrl), 0.6 ± 0.0 (12 wks, ctrl), p= <0.0001; feret: 20.1 ± 0.5 (P21, ctrl), 19.2 ± 0.5 (12 wks, ctrl), p= 0.1842; feret angle: 98.6 ± 3.9 (P21, ctrl), 92.0 ± 4.2 (12 wks, ctrl), p= 0.2477; minferet: 13.1 ± 0.3 (P21, ctrl), 11.6 ± 0.3 (12 wks, ctrl), p= <0.0001; roudness: 0.7 ± 0.0 (P21, ctrl), 0.7 ± 0.0 (12 wks, ctrl), p= 0.0050; solidity: 0.8 ± 0.0 (P21, ctrl), 0.8 ± 0.8 (12 wks, ctrl), p= <0.0001, all unpaired t test with Welch’s correction). Similar decreases in soma area, circularity, and solidity were observed in the Snap25 cKO brains; however, a decrease in the feret angle was also detected. (Figure 7c, e) (soma area: 147 ± 4.0 (P21, cKO), 129 ± 4.7 (12 wks, cKO), p= 0.0028; perimeter: 55.3 ± 1.4 (P21, cKO), 56.6 ± 2.1 (12 wks, cKO), p= 0.6048; circularity: 0.6 ± 0.0 (P21, cKO), 0.6 ± 0.0 (12 wks, cKO), p= 0.0020; feret: 19.6 ± 0.4 (P21, cKO), 19.4 ± 0.6 (12 wks, cKO), p= 0.8241; feret angle: 109 ± 3.3 (P21, cKO), 96.7 ± 4.1 (12 wks, cKO), p= 0.0198; minferet: 12.2 ± 0.2 (P21, cKO), 11.9 ± 0.3 (12 wks, cKO), p= 0.4639; roundness: 0.7 ± 0.0 (P21, cKO), 0.7 ± 0.0 (12 wks, cKO), p= 0.6391; solidity: 0.9 ± 0.0 (P21, cKO), 0.8 ± 0.0 (12 wks, cKO), p= <0.0001, all unpaired t test with Welch’s correction).

**Figure 7.**
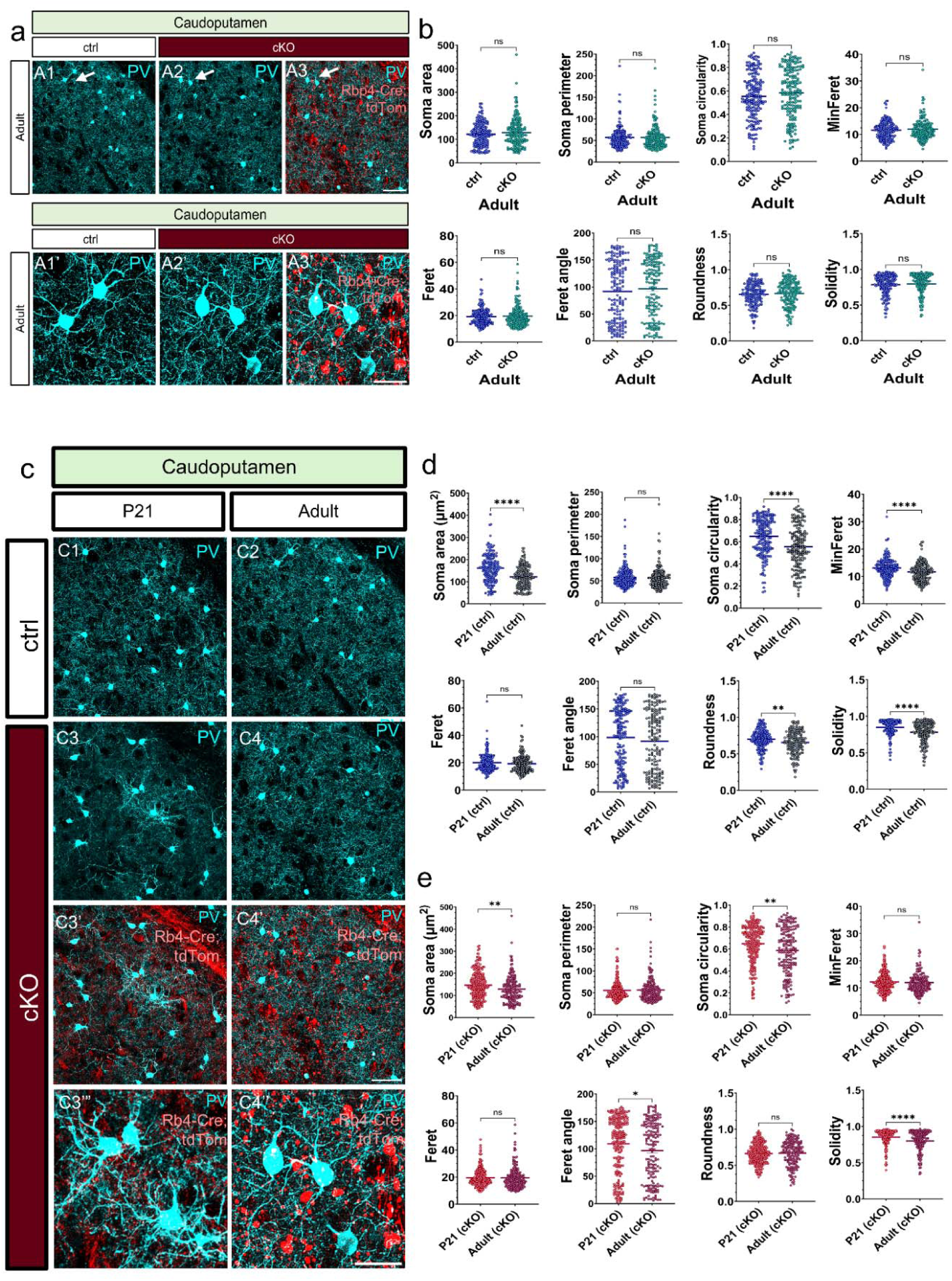
Chronic impairments in layer 5 activity have no permanent impact on the morphology of striatal interneurons in the adult brain. Maximum-intensity projected z-stack images of PV+ interneurons in the caudoputamen in the ctrl (A1, A1’) and the L5-silenced mice at 3 months of age (A2, A2’). Superimposed maximum-intensity projected z-stack confocal images of PV+ neurons and *Rbp4-Cre;Ai14;Snap25^fl/fl^* projection neurons in the striatum of adult Snap25 cKO mice (A3, A3’). White arrows mark the location of striatal PV+ neurons depicted in higher magnification (A1-A3’). No significant differences were observed in the soma parameters of PV+ neurons in the striatum of L5-silenced mice compared to control mice at 12 weeks of age (b). PV+ interneurons in the caudoputamen of ctrl mice at P21 (C1) vs 12 weeks of age (C2). PV+ interneurons in the caudoputamen of Snap25 cKO mice at P21 (C3) and 12 weeks of age (C4). Axonal projections of *Rbp4-Cre;Ai14;Snap25^fl/fl^* neurons in the caudoputamen in the L5-silenced mice at P21 (C3’, C3’’) and 12 weeks of age (C4’, C4”). Note the stark contrast in the axonal fibres of L5 projection neurons between the developing vs the adult Snap25 cKO mice. Accumulation of tdTomato+ punctae and fragmentation of L5 axons are only observed in the adult Snap25 cKO mice. Quantitative analysis of the trajectory of PV neuron morphology from P21 to 12 weeks of age disclosed a marked decrease in the soma area, circularity, minferet, roundness, and solidity of striatal PV neurons in the ctrl brains (d) (P21, n=180; 12 wks, n=168 cells, **** p<0.0001, unpaired t test with Welch’s correction). Similar reductions were noted in the soma area, circularity, feret angle, and solidity of striatal PV+ neurons in the L5-silenced mice between P21 and 12 weeks of age (e) (soma area, p= 0.0028, circularity, p= 0.0020, feret angle, p= 0.0198) (P21, n=236; 12 wks, n=171 cells). Scale bars: 50 µm (A3, C4’), 20 µm (A3’, C4’’).

These results demonstrate that chronic silencing of L5 activity modifies the morphological trajectory of striatal PV neurons as well as their soma characteristics at P21 while leaving their density intact. It is tempting to assume that the chronic modulation of Ca^2+^-dependent synaptic transmission in L5 has a temporary and transient effect on striatal PV+ neurons, given that the observed variations in striatal PV morphology were no longer detectable at twelve weeks of age. Please refer to Supplementary Table 4 for an overview of the morphometrics data.

### The absence of regulated vesicle release from Rbp4-Cre+ layer 5 does not alter the laminar organisation of PV interneurons in the juvenile brain

After establishing that the cessation of neurotransmitter release from L5 projection neurons from birth does not affect the density of cortical and subcortical PV neurons during development, we further investigated whether the chronic manipulation of L5 influenced the laminar distribution of PV neurons. No differences were detected in the laminar distribution of PV in M1 at P14 (Figure 8a A1-A2’, 8c) (P14 M1 L1: 0 ± 0 (ctrl), 0 ± 0 (cKO), p= >0.9999; L2/3: 106.8 ± 6.5 (ctrl), 164.7 ± 52.9 (cKO), p= 0.5340; L5: 129.3 ± 5.6 (ctrl), 174.7 ± 43.1 (cKO), p= 0.7345; L6: 59.0 ± 8.2 (ctrl), 87.1 ± 42.8 (cKO), p= 0.9370, two-way ANOVA with Šídák’s multiple comparisons test). The laminar distribution of PV neurons in M1 remained unchanged at P21 as well (Figure 8b B1-B2’, 8d) (P21 M1 L1: 0 ± 0 (ctrl), 0 ± 0 (cKO), p= >0.9999; L2/3: 126.8 ± 9.7 (ctrl), 150.3 ± 22.4 (cKO), p= 0.6876; L5: 145.2 ± 9.8 (ctrl), 170.0 ± 25.1 (cKO), p= 0.6454; L6: 38.8 ± 10.6 (ctrl), 68.7 ± 13.0 (cKO), p= 0.4757, two-way ANOVA with Šídák’s multiple comparisons test). We found that the absence of regulated vesicle release from L5 projection neurons did not alter the distribution of PV in any cortical layers in S1 at P14 (Figure 8g G1-G1’, 8i) (P14 S1 L1: 0 ± 0 (ctrl), 0 ± 0 (cKO), p= >0.9999; L2/3: 96.1 ± 19.0 (ctrl), 93.3 ± 29.2 (cKO), p= >0.9999; L4: 336.1 ± 49.5, 390.1 ± 130.4 (cKO), p= 0.9541; L5: 186.3 ± 18.2 (ctrl), 243.2 ± 63.0 (cKO), p= 0.9434; L6: 63.2 ± 5.6 (ctrl), 106.4 ± 26.9 (cKO), p= 0.9824, two-way ANOVA with Šídák’s multiple comparisons test). It can be concluded that Ca^2+-^dependent neurotransmission from layer 5 does not regulate the spatial organisation of PV neurons during postnatal development (Figure 8h H1-H1’, 8j) (P21 S1 L1: 0 ± 0 (ctrl), 0 ± 0 (cKO), p= >0.9999; L2/3: 122.3 ± 10.7 (ctrl), 124.8 ± 13.7 (cKO), p= >0.9999; L4: 352.6 ± 53.3 (ctrl), 307.1 ± 26.9 (cKO), p= 0.6483; L5: 190.7 ± 21.1 (ctrl), 214.5 ± 23.3 (cKO), p= 0.9640; L6: 68.8 ± 19.8 (ctrl), 99.4 ± 21.4 (cKO), p= 0.9024, two-way ANOVA with Šídák’s multiple comparisons test).

**Figure 8.**
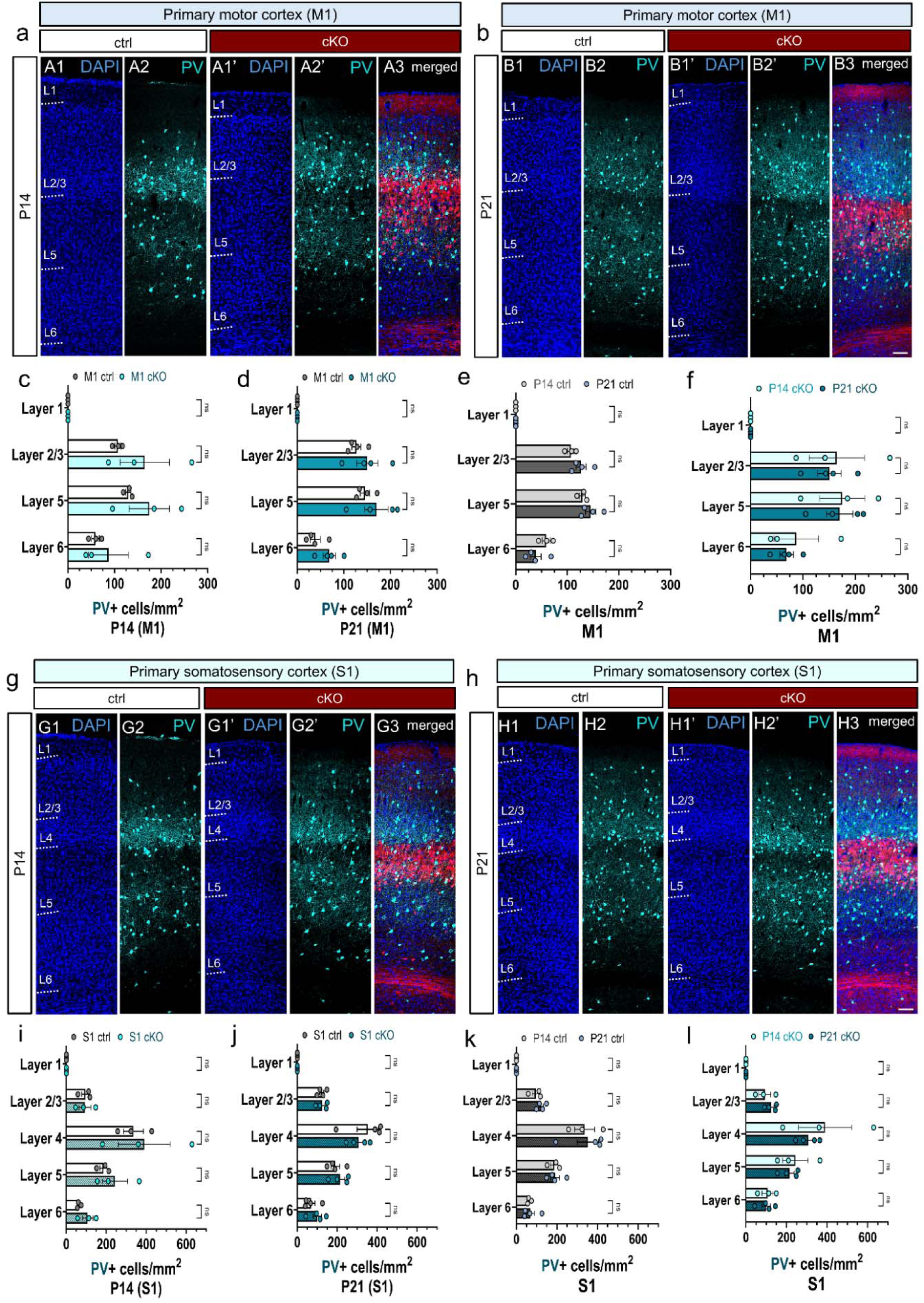
Chronic abolition of Ca^2+-^dependent neurotransmission from layer 5 pyramidal neurons does not disrupt the laminar organization of PV interneurons in the juvenile cortex. Laser-scanning confocal tile-scan images of PV+ immunostaining in the L5-silenced ctrl and cKO brains at P14 and P21. Nuclear counterstaining using DAPI was performed to define the boundaries of cortical layers in M1 (A1, A1’, B1, B1’) and S1 (G1, G1’, H1, H1’). Confocal tile-scans showing the distribution of PV+ neurons across the cortical layers in M1 at P14 (A2, A2’) and P21 (B2, B2’). Confocal tile scans depicting the distribution of PV+ neurons across the cortical layers in S1 at P14 (G2, G2’) and P21 (H2, H2’). Merged confocal images displaying *Rbp4-Cre;Ai14;Snap25^fl/fl^*projection neurons in red and PV-immunoreactive neurons in cyan in M1 (A3, B3) and S1 (G3, H3). Quantification of the laminar distribution of PV+ neurons in M1 at P14 and P21 (c, d). No significant differences were found in the density of PV neurons in any cortical layers in M1 between the ctrl and the L5-silenced brains at P14 and P21 (c, d). The developmental trajectory of the distribution of PV neurons in M1 in the L5-silenced ctrl brains did not reveal significant alterations in any cortical layers (e). The trajectory of the laminar distribution of PV neurons in M1 remained unaltered in the chronically silenced L5 brains between P14 and P21 (f). The distribution of PV neurons across the cortical layers in S1 showed no differences between P14 and P21 in the control and the L5-silenced brains (g-j). No differences were noted in the trajectory of the laminar distribution of PV+ neurons between P14 and P21 in the S1 ctrl and the L5-silenced brains (k, l). Scale bars: 100 µm. All data is represented as mean ± SEM values. 2-way ANOVA with Šídák’s multiple comparisons test. M1: primary motor cortex, S1: primary somatosensory cortex.

Having confirmed that there is no genotype effect on the laminar arrangement of PV interneurons in M1 and S1, we next sought to examine if developmental age affected the spatial organisation of PV. Between P14 and P21, we did not observe a significant change in the laminar distribution of PV neurons in the ctrl brains in M1 (Figure 8e) (M1 L1: 0 ± 0 (P14, ctrl), 0 ± 0 (P21, ctrl) p=>0.9999, L2/3: 106.8 ± 6.5 (P14, ctrl), 126.8 ± 9.7 (P21, ctrl), p=0.3337, L5: 129.3 ± 5.6 (P14, ctrl), 145.2 ± 9.9 (P21, ctrl), p=0.5500, L6: 59.0 ± 8.2 (P14, ctrl), 38.8 ± 10.6 (P21, ctrl), p=0.3260, two-way ANOVA with Šídák’s multiple comparisons test). The developmental trajectory of PV neurons in M1 in the L5-silenced brains was identical to that of the ctrl brain (Figure 8f) (M1 L1: : 0 ± 0 (P14, cKO), 0 ± 0 (P21, cKO), p= >0.9999, L2/3: 164.7 ± 52.9 (P14, cKO), 150.3 ± 22.4 (P21, cKO), p=0.9939, L5: 174.7 ± 43.1 (P14, cKO), 170.0 ± 25.1 (P21, cKO), p=>0.9999, L6: 87.1 ± 42.8 (P14, cKO), 68.7 ± 13.0 (P21, cKO), p=0.9849, two-way ANOVA with Šídák’s multiple comparisons test). Regarding the trajectory of PV interneurons distribution in S1 between P14 and P21, we observed no alterations in their laminar positioning in S1 in any cortical layers in the ctrl brains (Figure 8k) (S1 L1: 0 ± 0 (P14, ctrl), 0 ± 0 (P21, ctrl), p=>0.9999, L2/3: 96.1 ± 19.0 (P14, ctrl), 122.3 ± 10.7 (P21, ctrl), p=0.9706, L4: 336.1 ± 49.5 (P14, ctrl), 352.6 ± 53.3 (P21, ctrl), p=0.9964, L5: 186.3 ± 18.2 (P14, ctrl), 190.7 ± 21.1(P21, ctrl), p=>0.9999, L6: 63.2 ± 5.6 (P14, ctrl), 68.8 ± 19.8 (P21, ctrl), p=>0.9999, two-way ANOVA with Šídák’s multiple comparisons test). Silencing layer 5 had no impact on the trajectory of the laminar distribution of PV in S1 either (Figure 8l) (S1 L1: 0 ± 0 (P14, cKO), 0 ± 0 (P21, cKO), p=>0.9999, L2/3: 93.3 ± 29.2 (P14, cKO), 124.8 ± 13.7 (P21, cKO), p= 0.9909, L4: 390.1 ± 130.4 (P14, cKO), 307.1 ± 26.9 (P21, cKO), p=0.6379, L5: 243.2 ± 62.9 (P14, cKO), 214.5 ± 23.3 (P21, cKO), p= 0.9941, L6: 106.4 ± 26.9 (P14, cKO), 99.4 ± 21.4 (P21, cKO), p= >0.9999, two-way ANOVA with Šídák’s multiple comparisons test). These results indicate that the chronic manipulation of the activity of L5 neurons leaves the trajectory of PV distribution intact and does not rearrange the spatial organisation of PV during development. In agreement with the density results, we did not observe significant changes in the laminar density of PV neurons either in M1 or in S1 between P14 and P21.

### Chronic cessation of evoked vesicle release from L5 projection neurons does not influence the density of PV neurons in the adult cortex but disrupts the correlation between PV interneurons and the perineuronal net in M1

After verifying that the chronic abolition of Ca^2+-^dependent neurotransmission from L5 has no local or global effect on the density, laminar distribution, and trajectory of PV interneurons during the second and third week of postnatal development, we next explored whether chronically silencing L5 had a permanent impact on PV interneurons. To investigate whether different subsets of PV interneurons were differentially affected, we visualised the perineuronal net (PNN) using the *Vicia villosa* agglutinin (VVA) that allowed us to differentiate between PV neurons with PNN (PV+ VVA+) and PV neurons without PNN (PV+ VVA-). At three months of age, no differences were observed in the density of PV cells in M1 and S1 between the ctrl and the cKO brains even though the Rbp4-Cre;Snap25 cKO L5 neurons display clear signs of neurodegeneration and neuroinflammation by 3 months of age (Figure 9a A2, A2’, 9b B4’, 9c C2, C2’, 9d D4’) (see Hoerder-Suabedissen et al., 2019). Considering the drastic impairments of the silenced L5 projections, it is intriguing that cortical PV interneurons in M1 and S1 remain unaffected by the ongoing neurodegenerative events in the adult Snap25 cKO brains (Figure 9a A1, A1’, 9b B1, B1’, 9c C1, C1’, 9d D1, D1’, 9e) (M1: 109.3 ± 15.0 (ctrl), 105.0 ± 6.4 (cKO), p= 0.9700; S1: 132.5 ± 14.6 (ctrl), 161.0 ± 16.1 (cKO), p= 0.2887, two-way ANOVA with Šídák’s multiple comparisons test). To determine whether silencing L5 alters the PNN, we assessed the density of VVA+ cells in the adult cortex. There were no changes in the density of VVA+ cells in M1 and S1 between the ctrl and the cKO brains (Figure 9a A3, 9b B2, B2’, 9c C3, 9d D2, D2’, 9f) (M1: 147.7 ± 8.3 (ctrl), 112.1 ± 5.5 (cKO), p= 0.0544; S1: 220.4 ± 16.2 (ctrl), 250.8 ± 6.3 (cKO), p= 0.1044, two-way ANOVA with Šídák’s multiple comparisons test).

**Figure 9.**
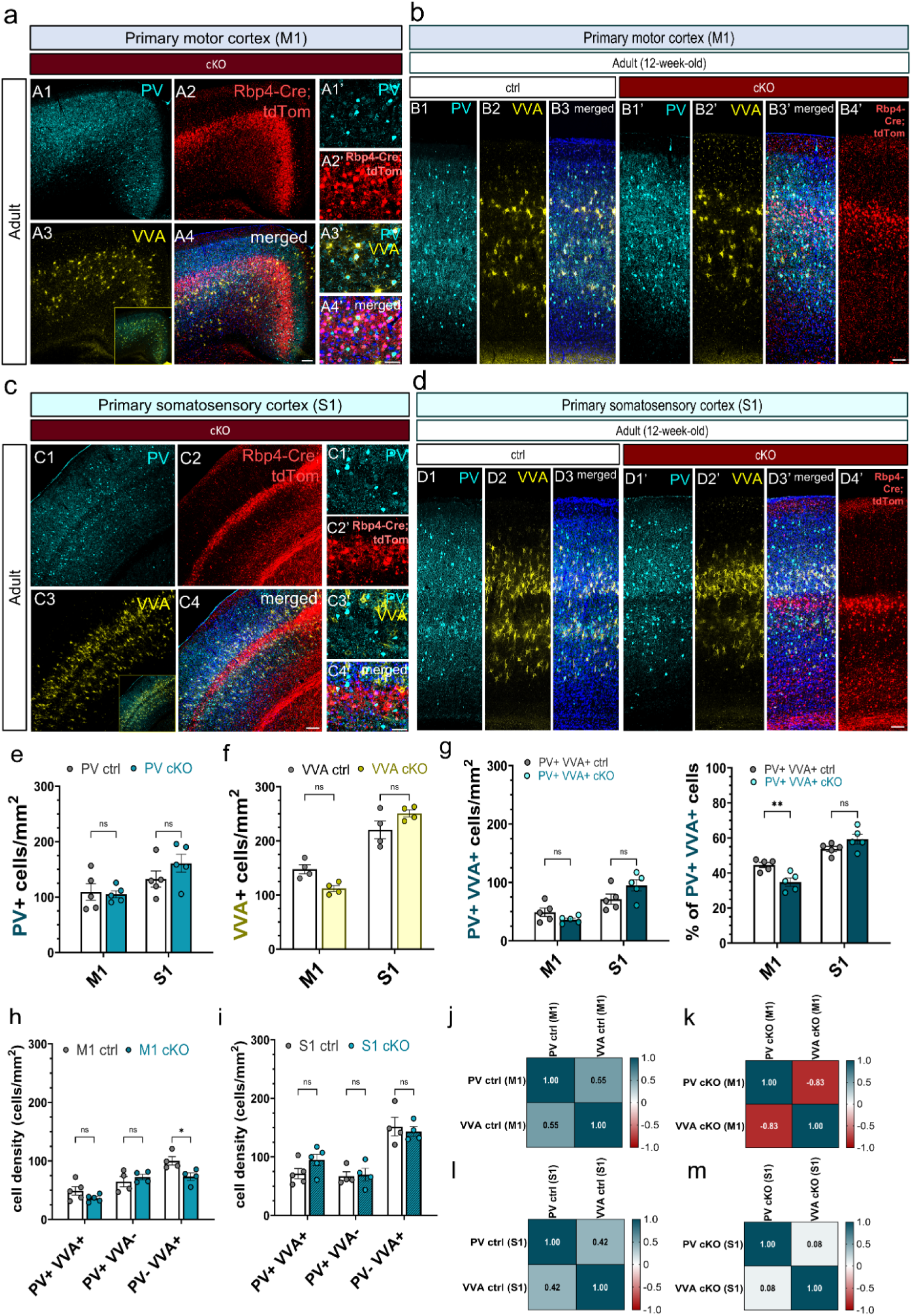
Long-term abolition of layer 5 pyramidal neuron activity alters the relationship between PV interneurons and the perineuronal net in M1, but it has little effect on the density of PV neurons in the adult brain. Low-magnification single-scan confocal images of the primary motor cortex in the Snap25 cKO brains at 3 months of age. PV-immunoreactive neurons are shown in cyan (A1, A1’), *Rbp4-Cre;Snap25^fl/fl^* projection neurons in L5 of M1 are seen in red (A2, A2’). The cells that are labelled with *Vicia Villosa* agglutinin (yellow) demonstrate the presence of perineuronal nets that encapsulate the somata of PV interneurons (A3, A3’). Merged confocal images show co-localisation of PV and VVA in M1 in Snap25 cKO brains. PV+ and VVA+ cells are highly concentrated near the cell bodies of L5 *Rbp4-Cre;Snap25^fl/fl^* projection neurons (A4, A4’). Maximum-intensity projected z-stack and confocal tile-scan images of M1 depicting PV-immunostained neurons (B1, B1’) and VVA-positive PNN labelling in the ctrl (B2) and L5-silenced brains at 12 weeks of age (B2’, B3). *Rbp4-Cre;Snap25^fl/fl^* projection neurons in the adult motor cortex showing signs of degeneration and fragmentation (B4’). Single-scan confocal images of PV+ neurons (C1, C1’) and VVA+ cells in the primary somatosensory cortex of Snap25 cKO brains at 3 months of age (C3, C3’). The degeneration of Rbp4-Cre+ neurons in layer 5 is a clear indication of the ongoing degenerative processes and heightened inflammation in Snap25 cKO mice (C2). Note the dying Rbp4-Cre;Snap25^fl/fl^ projection neurons at 3 months of age (C2’). PV+ and VVA+ signals superimposed on the *Rbp4-Cre,Ai14;Snap25^fl/fl^* neurons (C4, C4’). Combined tile-scan and maximum-intensity projected z-stack confocal images of PV-immunostained neurons (D1, D1’) and VVA-labelled PNNs in the ctrl and L5-silenced brains at 12 weeks of age (D2, D3, D2’, D3’). Degenerating Snap25-deleted Rbp4-Cre neurons in S1 in the adult Snap25 cKO mice. Note the accumulation of tdTomato-positive punctae throughout the cortex (D4’). Quantification of PV+ and VVA+ cells in M1 and S1 at 3 months of age revealed no significant differences in the density of PV+ and VVA+ neurons between the ctrl and the L5-silenced mice (e-f). The percentage of PV+ VVA+ cells in M1 was significantly lower in the Snap25 cKO mice at 3 months of age (p=0.0081, 2-way ANOVA with Šídák’s multiple comparisons test) (g). There was a significant decrease in the density of PV-VVA+ neurons in the adult motor cortex in the Snap25 cKO mice (h) (p= 0.0263, 2-way ANOVA with Šídák’s multiple comparisons test). Different subsets of PV neurons were not altered significantly in the adult S1 between the ctrl and the Snap25 cKO mice (i). PV-VVA correlation was impaired in M1 as the Snap25 cKO mice exhibited a negative correlation between PV and VVA (j-k) (r=-0.828). *p < 0.05, **p< 0.01. Scale bars: 100 µm (A4, C4), 50 µM (A4’, C4’, B3, D4). M1: primary motor cortex, S1: primary somatosensory cortex.

After confirming the abolition of evoked vesicle release had no long-term impact on the density of PV interneurons and the PNNs, we sought to explore changes in the subtypes of cortical PV interneurons. Three groups were distinguished based on the presence of PNN: 1) PV neuron with PNN or the double positives (PV+ VVA+); 2) PV neuron without PNN (PV+ VVA-); 3) PNN around non-PV neuron (PV-VVA+). While the density of the three subgroups (PV+ VVA+, PV+ VVA-, PV-VVA+) remained unchanged in S1 in the L5-silenced adult brains when compared to the ctrl brains (Figure 9i) (S1 PV+ VVA+: 71.4 ± 8.7 (ctrl), 94.9 ± 9.7(cKO), p=0.2707; PV+ VVA-: 67.0 ± 7.5 (ctrl), 69.5 ± 10.8 (cKO), p= 0.9978; PV-VVA+: 151.6 ± 15.8 (ctrl), 143.4 ± 8.1 (cKO), p= 0.9343, two-way ANOVA with Šídák’s multiple comparisons test); there was a significant decline in the density of the PV-VVA+ cells in M1 in the Snap25 cKO brains (Figure 9h) (M1 PV+ VVA+: 48.9 ± 7.2 (ctrl), 36.4 ± 2.7 (cKO), p= 0.3793; PV+ VVA-: 64.7 ± 8.8 (ctrl), 72.6 ± 4.6 (cKO), p= 0.7926; PV-VVA+: 100.2 ± 7.2 (ctrl), 73.2 ± 6.4 (cKO), p= 0.0263, two-way ANOVA with Šídák’s multiple comparisons test). It is notable that the decrease in the density of PV-VVA+ cells is happening without a change in the density of PV+ neurons indicating that the PNN is still plastic and capable of dynamic changes in the adult brain. We also noted a significant decrease in the percentage of the double positive cells (PV+ VVA+) in M1 in the layer 5-silenced mice (Figure 9g) (M1: 44.6% ± 1.4 (ctrl), 34.8% ± 2.3(cKO), p= 0.0081; S1: 53.7% ± 1.5 (ctrl), 59.3% ± 2.8 (cKO), p= 0.1479, two-way ANOVA with Šídák’s multiple comparisons test). Next, we explored how the chronic abolition of regulated vesicle release from L5 influenced the correlation between PV and VVA in different cortical regions. The PNN is thought to be predominantly surrounding PV interneurons, therefore the positive association between PV and VVA in the ctrl M1 and S1 areas is not an unanticipated result (Figure 9j, l). However, this correlation was disrupted in the motor cortex of L5-silenced mice (Figure 9k) (M1: r= 0.6 (ctrl), p=0.449; r= -0.8 (cKO), p=0.172). It is noteworthy that the effect of L5 appears to vary depending on the cortical region as, although decreasing, the correlation between PV and VVA in S1 did not reverse (Figure 9m) (S1: r=0.4 (ctrl), p=0.578; r=0.1 (cKO), p=0.923). The differential impact on M1 and S1 points to a variable degree of plasticity and correlation between PV and VVA in the adult cortices. For an overview of the adult and developmental PV and VVA results, please refer to Supplementary Table 5.

### Chronic silencing of layer 5 projection neurons has a differential and long-term effect on the laminar organisation of PNN and PV interneurons in the adult cortex

We then sought to study the laminar organisation of PV neurons and the PNN in the adult cortex. The chronic cessation of evoked vesicle release from L5 that leads to the degeneration of L5 neurons, and the disintegration of its long-range axonal projections may have permanent effects on the GABAergic interneurons that might not be apparent in the postnatal brain. At three months of age, we found a significant reduction in the density of the PNN in L5 and L2/3 of the motor cortex in the L5-silenced mice (Figure 10d) (M1 VVA+ L1: 0 ± 0 (ctrl), 0 ± 0 (cKO), p= >0.9999; L2/3: 205.1 ± 13.2 (ctrl), 154.2 ± 15.3 (cKO), p= 0.0028; L5: 214.6 ± 12.6 (ctrl), 159.2 ± 12.4 (cKO), p= 0.0011; L6a: 120.5 ± 10.8 (ctrl), 107.8 ± 5.1 (cKO), p= 0.8803; L6b: 2.0 ± 2.0 (ctrl), 0 ± 0 (cKO), p= >0.9999, two-way ANOVA with Šídák’s multiple comparisons test) and a similar trend was noted for L5 PV-VVA+ cells (Figure 10g) (M1 PV-VVA+ L1: 0 ± 0 (ctrl), 0 ± 0 (cKO), p= >0.9999; L2/3: 127.7 ± 11.5 (ctrl), 106.1 ± 19.0 (cKO), p= 0.5393; L5: 157.5 ± 11.5 (ctrl), 109.7 ± 6.6 (cKO), p= 0.0119; L6a: 81.1 ± 11.7 (ctrl), 71.2 ± 15.1 (cKO), p= 0.9670; L6b: 2.0 ± 2.0 (ctrl), 0 ± 0 (cKO), p= >0.9999, two-way ANOVA with Šídák’s multiple comparisons test). Of the subpopulations of PV neurons, only the PV+ VVA-subgroup showed a significant increase in L5 in the M1 region of the Snap25 cKO brains. This finding indicates that PV+ neurons without PNNs may re-enter a plastic phase by retaining their capacity to dynamically respond to changes in network activity even after the consolidation of the network and the closure of critical windows of development. (Figure 10a, b, c, e, f) (M1 PV+ VVA-L1: 0 ± 0 (ctrl), 0 ± 0 (cKO), p= >0.9999; L2/3: 77.1 ± 6.1 (ctrl), 81.9 ± 11.2 (cKO), p= 0.9984; L5: 82.1 ± 18.0 (ctrl), 131.1 ± 6.6 (cKO), p= 0.0052; L6a: 58.6 ± 9.2 (ctrl), 59.4 ±13.4 (cKO), p= >0.9999; L6b: 25.6 ± 9.9 (ctrl), 11.4 (cKO), p= 0.7992; M1 PV+ VVA+ L1: 0 ± 0 (ctrl), 0 ± 0 (cKO), p= >0.9999; L2/3: 72.5 ± 9.8 (ctrl), 48.8 ± 6.3 (cKO), p= 0.0562; L5: 56.6 ± 7.2 (ctrl), 56.6 ± 10.2 (cKO), p= >0.9999; L6a: 35.9 ± 5.7 (ctrl), 35.9 ± 7.9 (cKO), p= >0.9999; L6b: 3.8 ± 3.8 (ctrl), 0 ± 0 (cKO), p= 0.9962; M1 PV+ L1: 0 ± 0 (ctrl), 0 ± 0 (cKO); p= >0.9999; L2/3: 143.9 ± 15.8 (ctrl), 125.3 ± 15.2 (cKO), p= 0.8536; L5: 132.7 ± 21.6 (ctrl), 170.6 ± 13.1 (cKO), p= 0.2076; L6a: 93.2 ± 10.7 (ctrl), 91.3 ± 18.8 (cKO), p= >0.9999; L6b: 26.5 ± 7.7 (ctrl), 18.6 ± 6.5 (cKO), p= 0.9961, two-way ANOVA with Šídák’s multiple comparisons test). In contrast to M1, no differences were detected in the laminar distribution of VVA+ and PV-VVA+ cells in S1 in the L5-silenced brains (Figure 10k, n) (S1 VVA+ L1: 0 ± 0 (ctrl), 0 ± 0 (cKO), p= >0.9999; L2/3: 193.8 ± 14.4 (ctrl), 192.1 ± 9.9 (cKO), p= >0.9999; L4: 605.6 ± 61.9 (ctrl), 669.1 ± 27.4 (cKO), p= 0.2112; L5: 288.4 ± 7.9 (ctrl), 350.7 ± 16.4 (cKO), p= 0.2282; L6a: 164.7 ± 12.5 (ctrl), 192.7 ± 5.6 (cKO), p= 0.9252; L6b: 13.6 ± 2.6 (ctrl), 2.4 ± 2.4), p= 0.9994; S1 PV-VVA+ L1: 0 ± 0 (ctrl), 0 ± 0 (cKO); p= >0.9999; L2/3: 138.1 ± 10.2 (ctrl), 124.7 ± 13.4 (cKO), p= 0.9958; L4: 393.4 ± 43.1 (ctrl), 342.9 ± 27.3 (cKO), p= 0.2724; L5: 182.2 ± 8.0 (ctrl), 187.7 ± 24.4 (cKO), p= >0.9999; L6a: 118.5 ± 14.1 (ctrl), 129.0 ± 2.3 (cKO), p= 0.9989; L6b: 9.5 ± 4.0 (ctrl), 2.4 ± 2.4 (cKO), p= 0.9999, two-way ANOVA with Šídák’s multiple comparisons test). However, both the density of PV+ and PV+ VVA+ neurons rose significantly in L4 of S1 in the Snap25 cKO brains. (Figure 10j, l) (S1 PV+ L1: 0 ± 0 (ctrl), 0 ± 0 (cKO), p= >0.9999; L2/3: 110.4 ± 27.9 (ctrl), 150.2 ± 16.7(cKO), p= 0.6969; L4: 234.6 ± 36.2 (ctrl), 336.8 ± 34.3 (cKO), p= 0.0063; L5: 196.6 ± 16.6 (ctrl), 259.4 ± 28.4 (cKO), p= 0.2028; L6a: 97.9 ± 12.1 (ctrl), 110.5 ± 12.3 cKO), p= 0.9987; L6b: 48.8 ± 7.6 (ctrl), 42.4 ± 12.6 (cKO), p= >0.9999; S1 PV+ VVA+ L1: 0 ± 0 (ctrl), 0 ± 0 (cKO), p= >0.9999; L2/3: 53.4 ± 15.4 (ctrl), 60.9 ± 8.0 (cKO), p= 0.9999; L4: 188.5 ± 43.0 (ctrl), 289.9 ± 38.1 (cKO), p= 0.0023; L5: 100.3 ± 9.8 (ctrl), 147.5 ± 19.6 (cKO), p= 0.4017; L6a: 46.9 ± 8.4 (ctrl), 60.8 ± 6.1 cKO), p= 0.9961; L6b: 5.4 ± 6.2 (ctrl), 6.2 ± 6.2 (cKO), p= >0.9999; S1 PV+ VVA-L1: 0 ± 0 (ctrl), 0 ± 0 (cKO), p= >0.9999; L2/3: 74.4 ± 19.5 (ctrl), 88.7 ± 10.3 (cKO), p= 0.9740; L4: 48.1 ± 15.6 (ctrl), 31.8 ± 8.1 (cKO), p= 0.9510; L5: 93.2 ± 16.2 (ctrl), 113.8 ± 26.7 (cKO), p= 0.8651; L6a: 55.2 ± 4.9 (ctrl), 53.6 ± 10.1 (cKO), p= >0.9999; L6b: 51.9 ± 2.2 (ctrl), 39.8 ± 15.9 (cKO), p= 0.9886, two-way ANOVA with Šídák’s multiple comparisons test).

**Figure 10.**
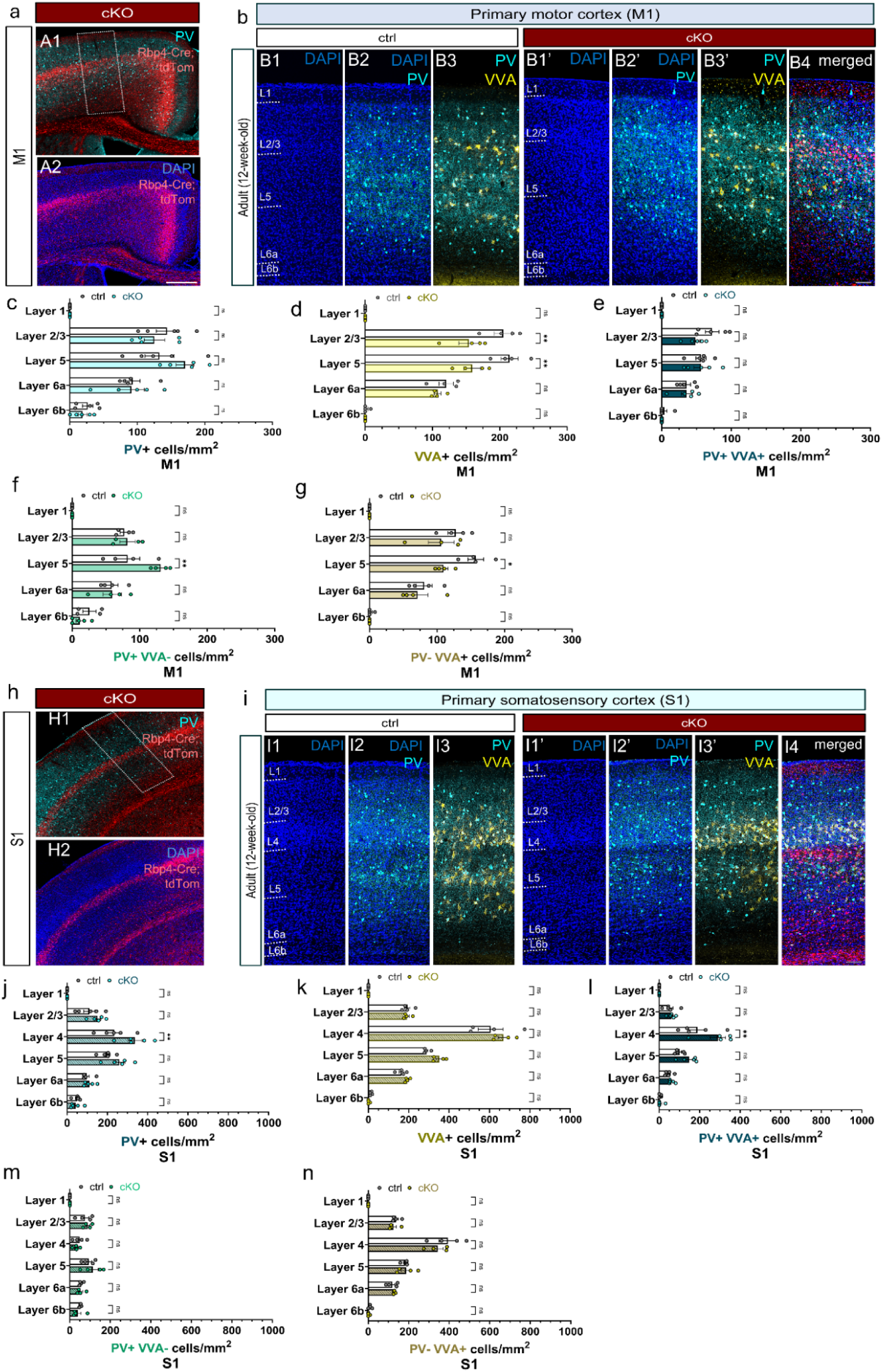
Chronic silencing of layer 5 projection neurons rearranges and differentially impacts the laminar distribution of PV interneurons and the perineuronal nets in the adult primary motor and somatosensory cortex. Low-magnification point-scanning confocal images of the primary motor and somatosensory cortices at 3 months of age illustrating PV interneurons (A1, H1) and L5-silenced projection neurons (A2, H2). High-magnification image of the white boxed region outlining M1 is shown in B4. Combined tile-scan and maximum-intensity projected z-stack confocal images representing the laminar distribution of PV+ and VVA+ cells in the ctrl and the Snap25 cKO mice in M1 (B1- B4) and S1 at 12 weeks of age (I1-I4). White dotted lines indicate the boundaries of each cortical layer. Laminar distribution analysis showed no significant changes in the density of PV+ and PV+ VVA+ in M1 at 12 weeks of age between the adult ctrl and the L5-silenced brains (c, e). The density of VVA+ cells was significantly reduced in the Snap25 cKO mice both in L2/3 and L5 in M1 (d) (L2/3, p= 0.0028; L5, p= 0.0011, 2-way ANOVA with Šídák’s multiple comparisons test). At three months of age, the Snap25 cKO motor cortex demonstrated a considerable rise in the density of PV+ VVA-cells in L5 and a significant decrease in the density of PV-VVA+ cells in L5 (f, g) (PV+ VVA-, p= 0.0052; PV- VVA+, p= 0.0119). Laminar distribution analysis of S1 revealed a significant increase in the density of PV+ neurons in L4 at 3 months of age in the L5-silenced mice (j) (p= 0.0063 ). The density of double positive PV+ VVA+ neurons followed the same trend in L4 in S1 (l) (p= 0.0023 ). The laminar distribution of VVA+, PV+ VVA-, and PV- VVA+ neurons was not altered significantly in any cortical layers in S1 at 3 months of age (k, m, n). Statistical tests: 2-way ANOVA with Šídák’s multiple comparisons. Scale bars: 200 µm (A2), 50 µm (B4, I4). M1: primary motor cortex, S1: primary somatosensory cortex.

Considering that the chronic abolition of synaptic vesicle release from L5 projection neurons has altered the laminar distribution of PV+ neurons and the PNNs only in the adult cortex, it is tempting to conclude that the effect of the chronic absence of neurotransmitter release from L5 on PV neurons is long-term and not manifested until adult stages. Since the total density of PV and VVA cells remained unchanged, the chronic silencing of L5 does not lead to interneuron cell death and loss of PNNs in the adult brain. However, alterations in the distribution of both PV and VVA on distinct cortical layers suggest a rearrangement of the laminar positioning of PV interneurons and the PNNs to compensate for the absence of synaptic transmission from L5.

### The correlation between PV and the perineuronal net is cortical layer-dependent

We previously established that the degree of correlation between PV and VVA neurons varied between different cortical regions and the chronic abolition of regulated vesicle release from layer 5 projection neurons disrupted the positive PV-VVA correlation in the adult primary motor cortex in the Snap25 cKO brains (Figure 8j,k). Considering that the correlation between PV and VVA can also be influenced by the laminar position of PV neurons and PNNs, we next examined the Pearson correlation between PV+ and VVA+ immunoreactive cells in different cortical layers of M1 and S1 at twelve weeks of age. We also sought to determine whether the association between PV and VVA cell density varied between the upper and lower cortical layers. In M1, the Snap25 cKO brains displayed a negative correlation between PV and VVA in L2/3 and L6a, in contrast to the ctrl brains’ positive correlation in L2/3 and L6a (Figure 11c) (M1 L2/3: r= 0.3 (ctrl), p=0.703; r= -0.4 (cKO), p=0.649; L5: 0.2 (ctrl), p=0.789; r= 0.4 (cKO), p=0.614; L6a: r=0.0 (ctrl), p=0.952; r= -0.4 (cKO), p=0.562; L6b: r= -0.5 (ctrl), p=0.451; r=0.0 (cKO)). In the L5-silenced brains, the correlation between PV and VVA in L2/3 and L4 of S1 was reversed (Figure 11d) (S1 L2/3: r= 0.0 (ctrl), p=0.984; r= -0.7 (cKO), p=0.328; L4: r= 0.0 (ctrl), p=0.962; r= -0.5 (cKO), p=0.493; L5: r=-0.9 (ctrl), p=0.075; r= -0.1 (cKO), p=0.892; L6a: r= 0.8 (ctrl), p=0.246; r= 0.6 (cKO), p=0.354; L6b: r= -0.5 (ctrl), p=0.467; r= -0.5 (cKO), p=0.525).

**Figure 11.**
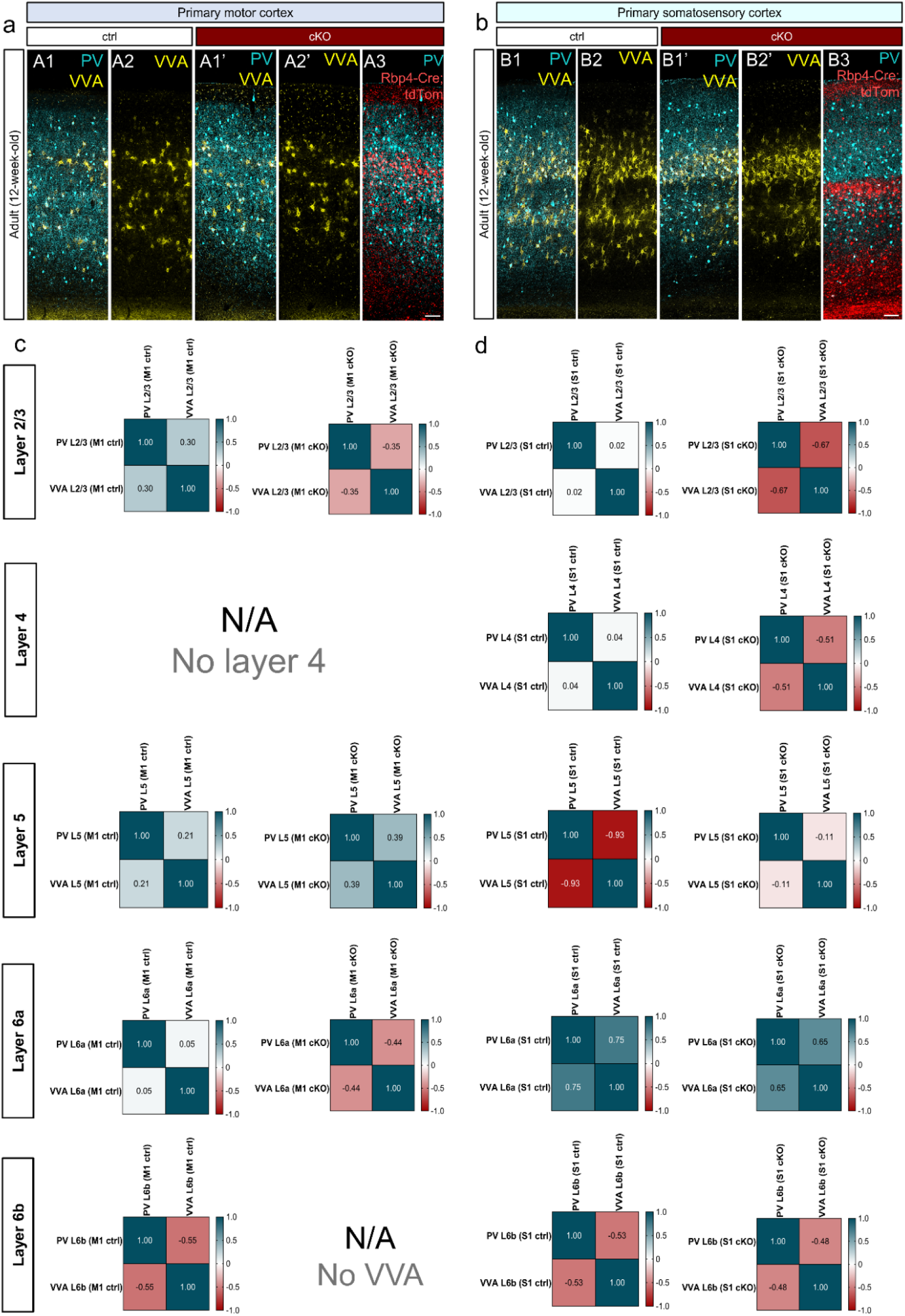
Chronic silencing of the activity of layer 5 projection neurons unveils regional and layer-dependent differences in the correlation between cortical PV interneurons and the perineuronal nets. Combined tile-scan and maximum-intensity projected z-stack confocal images depicting PV- immunostained neurons, VVA-labelled PNNs, and the degenerating *Rbp4-Cre;Ai14;Snap25^fl/fl^*projection neurons in M1 and S1 in the ctrl (A1, A2, B1, B2) and the chronically silenced L5 brains at 12 weeks of age (A1’-A3, B1’-B3). Heatmaps illustrating the Pearson correlation between PV+ and VVA+ cells in all cortical layers in M1 at 12 weeks of age (c). Note that the superficial and the infragranular cortical layers in M1 exhibit varying levels of correlation between PV+ and VVA+ in the ctrl and the Snap25 cKO brains. In the Snap25 cKO brains, there were no VVA+ cells in L6b in M1. The direction of correlation between PV+ and VVA+ was reversed in the L5-silenced mice in L2/3 and L6a of M1. Heatmaps illustrating the Pearson correlation between PV+ and VVA+ cells in all cortical layers in S1 at 12 weeks of age (d). Similar to M1, the correlation between PV and VVA in S1 varies greatly across cortical layers. Note that both the ctrl and the Snap25 cKO mice show a negative correlation in L6b between PV and VVA at 12 weeks of age. Numbers in heatmaps represent r values of Pearson correlation. Scale bars: 50 µm (A3, B3). M1: primary motor cortex, S1: primary somatosensory cortex.

The negative correlation between PV and VVA in the Snap25 cKO mice indicates that not all PV+ cells are surrounded by PNNs and while most PV+ neurons are VVA+ in L4 of S1, there are PNNs around neurons that are PV-. Although the correlation remained unaltered in L6a and L6b in the Snap25 cKO brains, the negative correlation in L6b observed both in the ctrl and the cKO brains supports our density data showing that VVA rarely occurs in L6b. The findings imply that the previously documented robust correlation between PV and VVA changes in degree throughout cortical layers and between the motor and somatosensory cortices. These results also demonstrate that upper and lower cortical layers in functionally distinct cortical areas have different correlation values. Establishing that PV-VVA correlation is both region-and layer dependent further highlights how dynamically PV and PNNs interact, and how their link is more nuanced than previously believed.

### Persistently silencing subsets of layer 5 projection neurons does not lead to long-term changes in the number of subcortical PV neurons in the projection regions of layer 5 in the adult brain

To investigate the long-term influence of the chronic cessation of evoked vesicle release on PV interneurons, we analysed the density of subcortical PV neurons in the output regions of L5 neurons at three months of age. It is important to note that at this age, most subcortical axons from the silenced L5 projection neurons have degenerated. Surprisingly, the layer 5-silenced brains did not reveal significant alterations in the density of subcortical PV neurons in any of the long-range projection regions of L5 (Figure 12a, 12c) (CPu: 64.3 ± 11.9 (ctrl), 65.8 ± 5.4 (cKO), p= >0.9999; GPe: 180.8 ± 19.3 (ctrl), 194.3 ± 14.9 (cKO), p= 0.9989; LP: 24.0 ± 6.3 (ctrl), 24.5 ± 3.2 (cKO), p= >0.9999; MD: 15.7 ± 4.0 (ctrl), 23.0 ± 5.5 (cKO), p= >0.9999; TRN: 765.0 ± 44.7 (ctrl), 746.3 ± 47.8 (cKO), p= 0.9966; SC: 341.0 ± 21.3 (ctrl), 327.3 ± 45.0 (cKO), p= 0.9989, two-way ANOVA with Šídák’s multiple comparisons test). In the striatum, neither the density of the PNN nor the density of PV- VVA+ cells changed in the adult Snap25 cKO brains (Figure 12b, f, g) (CPu VVA+: 61.1 ± 3.1 (ctrl), 50.4 ± 7.1 (cKO), p= 0.2743; PV- VVA+: 25.2 ± 5.6 (ctrl), 21.4 ± 2.9 (cKO), p=0.8362; CPu PV+ VVA+: 55.2% ± 5.5 (ctrl), 44.9% ± 8.3 (cKO), p=0.5326, two-way ANOVA with Šídák’s multiple comparisons test). To determine if chronically abolishing neurotransmitter release from L5 changes the maturation of PV neurons into adulthood, we next looked at the trajectory of subcortical PV neurons from development to adulthood.

**Figure 12.**
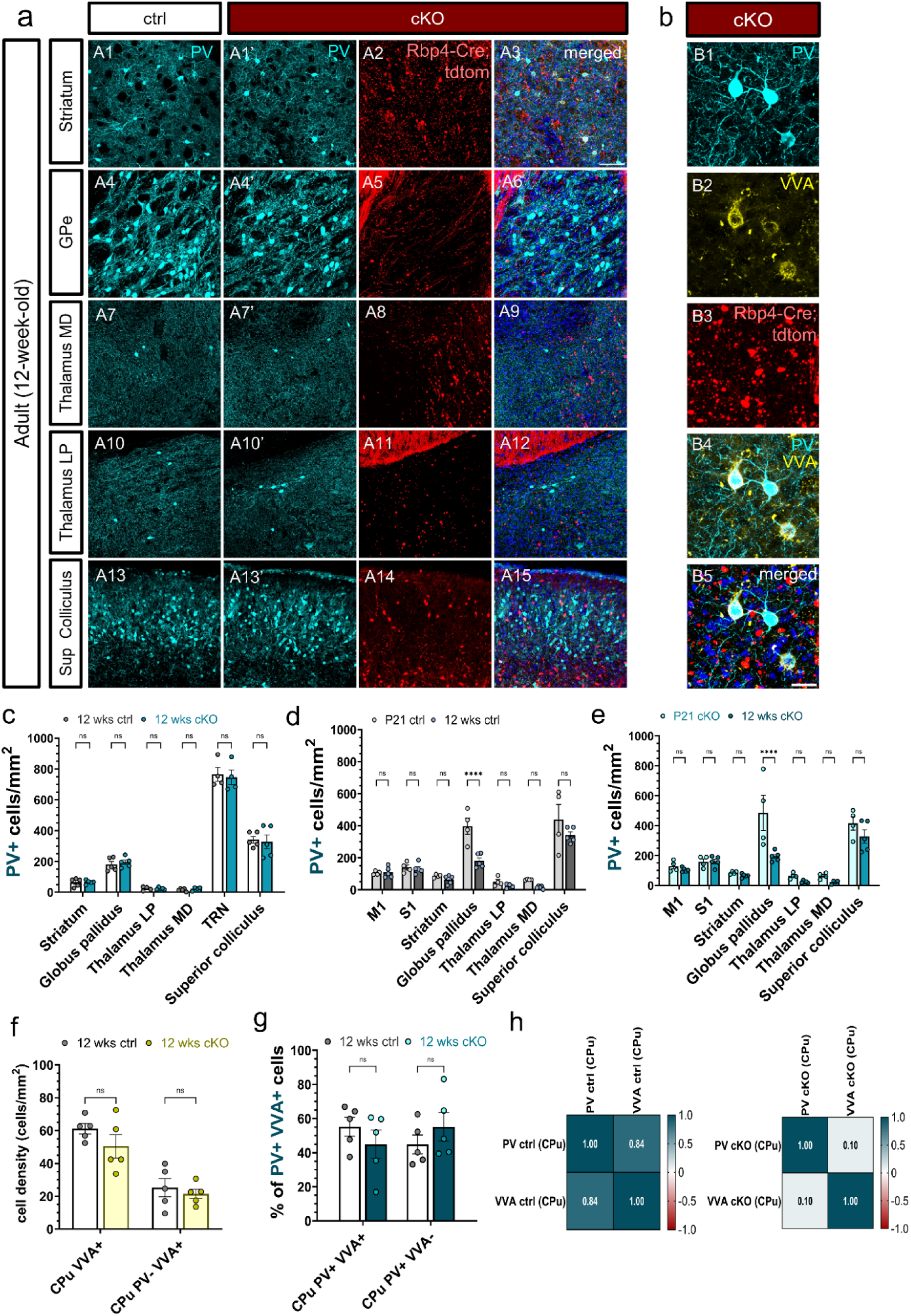
Loss of synaptic vesicle release from Rbp4-Cre+ layer 5 projection neurons has no long-term impact on the density of PV interneurons in the subcortical projection regions of layer 5. Maximum-intensity projected confocal z-stack images demonstrating the selected output regions of the silenced Rbp4-Cre projection neurons at 3 months of age. Subcortical PV+ neurons in the output regions of L5 are shown in cyan, while the *Rbp4-Cre;Ai14;Snap25^fl/fl^* are depicted in red. PV-immunopositive neurons in the caudoputamen and the external segment of the globus pallidus in the ctrl (A1, A4) and the Snap25 cKO mice (A1’, A4’). PV+ neurons in the higher-order lateral posterior and mediodorsal thalamic nuclei in the ctrl (A7, A10) and the Snap25 cKO mice (A7’, A10’). PV-immunoreactive neurons in the superior colliculus at 12 weeks of age in the ctrl (A13) and the L5-silenced mice (A13’). Long-range corticofugal projections of *Rbp4- Cre;Ai14;Snap25^fl/fl^* cortical pyramidal neurons in the adult Snap25 cKO mice in the basal ganglia (A2-A6), the higher-order thalamic nuclei (A8-A12), and the midbrain (A14-A15). Note the disintegration of the projections and the enlarged tdTomato+ swellings on the axonal fragments of *Rbp4-Cre;Ai14;Snap25^fl/fl^*neurons. Point-scanning maximum-intensity projected z-stack images of striatal PV+ neurons and VVA-labelled PNNs at 12 weeks of age in the Snap25 cKO mice (B1, B2). Disintegrating axonal fibres of the silenced L5 projection neurons in the caudoputamen (B3). Presence of perineuronal nets (PNNs) surrounding striatal PV+ neurons in adult L5-silenced mice (B4, B5). Quantification of PV+ neurons in the output regions of *Rbp4- Cre;Ai14;Snap25^fl/fl^* neurons at 12 weeks of age. No subcortical regions showed statistically significant variations in PV+ neuron density between the Snap25 cKO and control groups (c). Both in the ctrl and the Snap25 cKO mice, the trajectory of PV+ neurons between P21 and 3 months of age showed a marked decrease in PV+ neuron density in the globus pallidus (p****<0.0001, 2-way ANOVA with Šídák’s multiple comparisons test) (d, e). There were no significant variations in the density of VVA+ and PV- VVA+ cells in the adult striatum (f). In the striatum, the layer 5-silenced animals exhibited a reduction in the degree of positive association between PV and VVA (h). Scale bars: 50 µm.

First, we examined the course of maturation of subcortical PV neurons between P21 and 3 months of age in the ctrl brains where we noted a significant decrease in PV cell density in the GPe (Figure 12d) (P21: 397.0 ± 50.8 (ctrl), 12 wks: 180.8 ± 19.3 (ctrl), p= <0.0001, two-way ANOVA with Šídák’s multiple comparisons test). We noticed the same trend in the layer 5- silenced brains. Of the output regions of L5, the density of PV neurons was only altered in the GPe, while the rest of the projection sites and the cortical regions remained unchanged (Figure 12e) (M1: 129.0 ± 17.8 (P21 cKO), 105.0 ± 6.4(12 wks cKO), p= 0.9990; S1: 158.3 ± 19.2 (P21 cKO), 160.5 ± 15.9 (12 wks cKO), p= >0.9999; CPu: 85.3 ± 4.9 (P21 cKO), 65.8 ± 5.4 (12 wks cKO), p= 0.9997; GPe: 485.0 ± 117.4 (P21 cKO), 194.3 ± 14.9 (12 wks cKO), p= <0.0001; LP: 63.6 ± 10.0 (P21 cKO), 24.5 ± 3.2 (12 wks cKO), p= 0.9796; MD: 64.3 ± 10.2 (P21 cKO), 23.0 ± 5.5 (12 wks cKO), p= 0.9794; 415.7 ± 45.9 (P21 cKO), 327.4 ± 45.0 (12 wks cKO), p= 0.4268, two-way ANOVA with Šídák’s multiple comparisons test). These findings are consistent with the idea that the first week of development marks the most significant milestones in the maturation of PV neurons, and when these neurons undergo developmental apoptosis, their populations appear to stabilise and hold steady into adulthood (Southwell et al., 2012; Denaxa et al., 2018; Wong et al., 2018, 2022; Magno et al; 2021). Having established that the chronic abolition of vesicle release from L5 perturbed the correlation between PV and VVA in the adult motor cortex, we next examined whether PV density correlated with VVA density in subcortical regions. Although the striatum of the layer 5-silenced mice showed a positive PV-VVA correlation (Figure 12h), the level of correlation was not as great as that of the ctrl brains (CPu: r=0.8 (ctrl), p=0.075; r=0.1 (cKO), p=0.879). These results suggest that the chronic abolition of Ca2+- dependent neurotransmission from L5 neurons has a differential impact on cortical and subcortical brain regions, and the correlation between PV and VVA may be region-specific.

## Discussion

We explored the role of deep-layer projection neurons in the spatial and laminar organisation of cortical and subcortical parvalbumin neurons at distinct developmental stages and in adulthood. Using a triple transgenic conditional knockout mouse (*Rbp4-Cre;Ai14;Snap25^fl/fl^*), we selectively ablated *SNAP25* from Rbp4-Cre-expressing neurons that abolished evoked synaptic vesicle release from subsets of layer 5 corticostriatal pyramidal neurons (Welch et al., 2000; Marques- Smith et al., 2016; Gustus et al., 2018; Hoerder-Suabedissen et al., 2019; Korrell et al., 2019, Hayashi et al., 2021; Krone et al., 2021). We tested the hypothesis that infragranular pyramidal cells instruct the organisation of inhibitory cortical circuits by fine-tuning the number of GABAergic cells via activity-dependent mechanisms. We found that perturbing the function of infragranular pyramidal cells by abolishing Ca^2+^-dependent neurotransmission from L5 did not impede the development of PV+ neurons; however, it rearranged the laminar distribution of PV+ cells in the adult motor and somatosensory cortex. We demonstrated that chronic silencing of L5 had a differential impact on PV neurons and the PNNs by affecting different cortical layers. While the absence of vesicle release from L5 impacted PV+ neurons in L4 in S1, it altered the laminar distribution of VVA+ and PV- VVA+ neurons in L2/3 and L5 in M1. These results demonstrate that different cortical layers and distinct subtypes of PV+ neurons respond differently to the abolition of vesicle release from L5 projection neurons. We also discovered that the correlation between PV and VVA varies by cortical regions and layers and such correlation can be disrupted in the adult brain by chronically manipulating L5 projection neurons.

Recent research has demonstrated that the bidirectional manipulation of pyramidal neuron activity using Designer Receptors Exclusively Activated by Designer Drugs (DREADDs) only elicited a change in the density of PV+ and SST+ interneurons if the chemogenetic manipulation was performed between P5 and P8 (Wong et al., 2018, 2022). Considering that no such effect was detected when the pyramidal cell activation/inhibition occurred after the developmental apoptosis of GABAergic cells between P10 and P13, it appears there is a critical window of development when the pyramidal neuron activity is indispensable for the survival of GABAergic neurons. Developmental interneuron apoptosis is reported to occur between P5 and P10 with a peak at P7 (Wong et al., 2018; Southwell et al., 2012; Lim et al., 2018; Pfisterer and Khodosevich, 2017); therefore, most studies restricted the manipulation of pyramidal neuron activity to the window of programmed cell death (Wong et al., 2018; 2022; Sreenivasan et al., 2022) or carried out embryonic and/or neonatal interventions (Priya et al., 2018; Denaxa et al., 2018; Duan et al., 2020). In our study, pyramidal cell activity is irreversibly and chronically silenced in the *Rbp4-Cre;Ai14;Snap25^fl/fl^* mice and thus, our results cannot directly be compared to the results of studies using transient modulation of pyramidal cell activity (days vs lifelong manipulation). However, the mixed outcomes of pyramidal neuron activity on interneuron density reveal that GABAergic cells have different degrees of vulnerability during development highlighting the importance of studying GABAergic neurons at distinct stages of development.

Our work also investigated the global effects of chronically silencing the activity of glutamatergic L5 projection neurons and revealed no significant alterations in the density of subcortical PV+ interneurons in the long-range projection targets of Rbp4-Cre+ pyramidal neurons. Previous research has highlighted the role of cortical control in the regulation of striatal PV+ interneurons in the developing brain and argued that the cortical inputs determine the ultimate number of interneurons (Sreenivasan et al., 2022). Our study has different findings since the density of PV+ immunoreactive neurons in the striatum remained unchanged at P14 and P21 following the chronic ablation of Snap25 from L5 projection neurons. While we detected changes in the morphology of striatal PV interneurons at P21; however, these changes diminished in the adult brain suggesting that the cessation of neurotransmitter release from L5 only had a transient effect on PV morphology. Surprisingly, the developmental trajectory of PV morphology was altered in the layer 5-silenced mice.

The differences in findings regarding the density of striatal PV neurons may be explained by variations in the methods used to manipulate pyramidal neuron activity and the temporal differences in these manipulations. Sreenivasan and colleagues performed DREADD injections to carry out an acute and reversible manipulation of pyramidal neuron activity for days. As opposed to our research, which sought to detect long-term alterations in the distribution of PV neurons by employing a chronic and irreversible modification of L5 pyramidal neurons. Another interesting question related to the role of corticostriatal pyramidal neurons in establishing the final number of interneurons both in cortical and subcortical circuits is the conditional deletion of different SNARE proteins from L5 projection neurons. Our results showed that the genetic ablation of *SNAP25* from subpopulations of Rbp4-Cre-expressing L5 projection neurons left the density and laminar distribution of PV interneurons intact during the early stages of development both in the cortex and the subcortical innervation sites of L5. This contradicts the results of earlier studies where another fusion protein of the SNARE machinery, i.e., syntaxin-binding protein 1 (*Syt1*), was genetically ablated from Rbp4-Cre+ neurons and caused a reduction in the number of PV neurons in the striatum (Sreenivasan et al., 2022).

The question arises as to whether the loss of different presynaptic SNARE fusion proteins has a differential impact on PV cell number. Our results suggest that Snap25 and Ca^2+^- dependent synaptic transmission is dispensable for PV survival given the lack of local and global effect of abolished vesicle release from subgroups of cortical L5 projection neurons. However, in agreement with other studies (Sreenivasan et al., 2022), the conditional ablation of Munc18-1 from Rbp4-Cre+ L5 neurons only had a subtle effect on striatal PV density. This is an interesting observation considering that the genetic removal of Munc18-1 results in the total cessation of neurotransmitter secretion (both evoked and spontaneous vesicle release are abolished) (Verhage et al., 2000). The ablation of Snap25 in our mouse model only affected regulated synaptic exocytosis and it is tempting to speculate that more severe perturbations in synaptic transmission would lead to a more severe phenotype. However, one probable explanation for the lack of drastic changes in the density and distribution of PV interneurons might be that the deletion of the presynaptic SNARE proteins only affects a portion of L5 pyramidal neurons. We have previously documented that 15% of NeuN+ cells are tagged by Rbp4-Cre::tdTomato (Hoerder-Suabedissen et al., 2019) highlighting that the majority of L5 glutamatergic neurons are unaffected by the ablation of Snap25. Considering that the rest of the non-Rbp4-Cre+ pyramidal neurons remain capable of regulated vesicle release, we speculate that these L5 pyramidal neurons may compensate for the cessation of activity of Rbp4-Cre+ L5 projection neurons. Moreover, we have not observed a significant loss in the density of the cell bodies of Rbp4-Cre+ neurons at P21, and a notable decline in Rbp4-Cre+ neurons was only noted at 8 months of age (Hoerder-Suabedissen et al., 2019). This might explain why PV interneurons survive the chronic abolition of synaptic transmission and remain unaffected in cortical and subcortical regions as well.

Cortical layer 5 and a subset of Drd1a-Cre-expressing layer 6b neurons have unique reciprocal connections with the thalamus and both deep-layer projection neurons selectively innervate the higher-order thalamic nuclei (Mo and Sherman, 2018; Hoerder-Suabedissen et al., 2019; Hayashi et al., 2021; Casas-Torremocha et al., 2022; Zolnik et al., 2023). The lateral posterior nucleus of the thalamus (LP) is one of the higher-order thalamic nuclei innervated by L5 axon terminals and therefore, appeared to be a good candidate region for assessing the global effect of silencing L5 on the density of PV+ neurons. But neither in the developing nor in the adult brain did PV+ neurons in LP or the mediodorsal (MD) nucleus of the thalamus exhibit significant changes in their density. The spatial distribution of PV may be defined by its input areas and the strength of these inputs. Regarding the thalamus, the ventral posteromedial nucleus of the thalamus (VPm) is found to be the single biggest source of input for the three canonical interneuron types (Wall et al., 2016; Hafner et al., 2019). However, the scarce number of PV neurons present in the rodent thalamus makes the investigation of the interaction between the thalamic projections of L5 and PV challenging.

We also assessed the long-term effects of silencing L5 on the density of cortical and subcortical PV neurons at 3 months of age. Consistent with the developmental results, the density and morphology of PV interneurons remained unaffected in the adult layer 5-silenced mice in all the cortical and subcortical regions investigated. The lack of effect of the chronic manipulation of L5 on PV density in the adult brain indicates that alterations in PV cell number are highly unlikely once the final number of GABAergic interneurons is determined during early development. However, their laminar positioning can still be influenced by chronically perturbing L5 pyramidal neurons. Since parvalbumin is activity-dependent (Patz et al., 2004), we would argue that density changes are not driven by cell loss, but by a dynamic up-or downregulation of the parvalbumin protein that is responding to changes in the cortical network and changes in pyramidal neuron activity. This is supported by recent research showing that the parvalbumin-expressing GABA interneurons are not eliminated in a mouse model of autism spectrum disorder and the downregulation of the parvalbumin protein expression is the cause of the apparent cell loss (Filice et al., 2016). The results of our recent study agree with this given that no major alterations were noted in the density of PV+ and VVA+ cells following the acute chemogenetic manipulation of Rbp4-Cre+ neurons (Vadisiute et al., unpublished).

Previous studies have demonstrated that fate-converting subcerebral projection neurons to callosal projection neurons altered the laminar distribution of GABAergic interneurons pointing to the role of glutamatergic pyramidal neuron identity in instructing the laminar arrangement of inhibitory neurons (Lodato et al., 2011; Wester et al., 2019). In our study, the chronic abolition of regulated vesicle release from L5 projection neurons changed the distribution of PV neurons in M1 and S1 at 3 months of age. We found a significant increase in the density of PV neurons in layer 4 of the primary somatosensory cortex. We also distinguished several subpopulations of interneurons given that there were PV cells that were not enwrapped by PNNs (PV+ VVA-) and there were PNNs that did not colocalise with PV (PV- VVA+). It has been proposed that PV interneurons may orchestrate the organisation and the state of PNNs surrounding them as transcriptomic analysis revealed that many of the necessary PNN lecticans and linkers, as well as membrane and secretory proteases, are expressed by PV neurons (Devienne et al., 2021; Dityatev et al., 2007, 2010; Ferrer and Dityatev, 2018; Kwok et al., 2012). The various subtypes of PV neurons (PV+ VVA+, PV+ VVA-, PV- VVA+) might reflect the individual response of PV neurons to pyramidal neuron activity as they up or downregulate their PNNs depending on the network state. It is probable that the PV+ VVA- subtype is more plastic and so retains its ability to change to network disturbances while the PV+ VVA+ subtype has limited plasticity as it is consolidated by PNNs. Therefore, the PNNs may endow PV neurons with distinct properties depending on whether they are surrounded by proteoglycans or not. Previous studies have discovered that through its control on potassium channel localization and synaptic AMPA receptor levels, brevican alters the intrinsic characteristics of PV+ interneurons and modulates PV function and excitability (Favuzzi et al., 2017).

The significant increase in the density of PV neurons in layer 4 of S1 at 3 months of age may indicate that the chronic cessation of L5 activity has a permanent effect on PV lamination that only manifests in adulthood. The concomitant increase in the density of PV+ VVA+ neurons in L4 may also suggest that PV interneurons try to counterbalance the absence of pyramidal neuron activity by upregulating the perineuronal net expression around them. Previous studies provide evidence that there is a direct correlation between PNN expression and thalamic innervation in primary sensory cortical regions (Lupori et al., 2023). It has also been shown that PNNs have the capacity to regulate thalamic afferents onto PV neurons and modulate visual input by thalamic recruitment of cortical PV interneurons (Faini et al., 2018). Our results show that chronically silencing L5 led to a significant decrease in the density of VVA+ and PV- VVA+ cells in L5 of M1 at 3 months of age. The alterations in the laminar arrangement of VVA+ neurons imply that the PNNs can be modulated by activity-dependent mechanisms and Ca^2+-^ dependent synaptic transmission may control PNN density in adult networks. The observed reductions in VVA+ in L5 of M1 may imply that adult cortical circuits are more receptive to high circuit plasticity when PNNs are decreased. The different patterns in the alterations of the laminar density of PV+ and VVA+ cells suggest that the influence of pyramidal cell activity on GABAergic interneurons and PNNs is regional and subtype-specific. Our results further support the regional and subtype-specific differences in interneuron density reported by previous studies (Duan et al., 2020; De Marco García et al., 2011; Ueno et al., 2018; Pouchelon et al., 2021).

Remarkably, we observed that the correlation between PV and VVA was disrupted in the adult Snap25 cKO mice in M1. Layer 2/3 and layer 5 in M1 had a reverse correlation between PV and VVA in the Snap25-ablated mice. We also found that the PV and PNN correlation is region- and layer-dependent. There was a high degree of correlation between PV and VVA density in layer 6a in S1. However, cortical layer 5 and layer 6b of S1 revealed a negative correlation between PV and VVA density. These findings support prior studies that demonstrate the association between PV and PNN is extremely region-dependent, fluctuates throughout layers, and differs significantly between cortical areas (Lupori et al., 2023).

One of the major limitations of the present study is that the Rbp4-Cre is a pan L5 line and therefore, it is not truly restrictive to a specific subset of L5 pyramidal neurons. Since the Rbp4- Cre line is unable to differentiate between IT vs ET-type L5 projection neurons, other transgenic models with more precise genetic targeting of L5 pyramidal neuronal subgroups will be needed. Another limiting factor is the chronic manipulation of pyramidal neuron activity. The Rbp4- Cre;Snap25 cKO mice only permit the permanent and irreversible manipulation of pyramidal neuron activity. Engineering of new CreERT mouse lines will allow us to manipulate pyramidal neurons in an acute fashion and perturb pyramidal neuron function with temporal precision. While the axonal projections of Rbp4-Cre+ neurons remained intact in the Snap25 cKO mice until P21, severe neurodegeneration was detected in the adult brain. This makes it more difficult to distinguish between the effects of persistently silencing L5 projection neurons on PV interneurons and the impacts of cell death and inflammation, which may have an indirect impact on PV density, rather than the absence of vesicle release from L5.

In this study, we performed a chronic, irreversible manipulation of subsets of L5 projection neurons to unveil the role of deep-layer pyramidal neurons in instructing the laminar and spatial distribution of PV neurons. We demonstrated the absence of chronic vesicle release from L5 projection neurons does not disrupt the development of PV+ neurons, but it leads to a rearrangement of PV+ and VVA+ neurons in a region-and layer-specific manner. It appears that distinct subtypes of PV neurons rely on different survival signals and region-specific mechanisms govern PV function and number. Further studies examining PV and PNN interactions will uniquely inform us about the plasticity of cortical circuits, the differential survival capacity of inhibitory interneurons, and the pathophysiological changes underpinning neuropsychiatric disorders with interneuronopathies and synaptopathies.

## Acknowledgements

We wish to thank all members of the Molnár lab for their stimulating discussions and critical reading of the manuscript. We are grateful to the Anatomical Society for funding this study and the DPhil studies of F.SZ. who held a PhD research studentship from the Anatomical Society (project title: ‘*The role of calcium-dependent neurotransmission in selected populations of cortical projection neurons in the distribution of GABAergic interneurons’*). Z.M.’s laboratory was supported by the following funding sources: BBSRC Project Grant (BB/X008711/1) – Brain mechanisms of sleep: top-down or bottom-up, with Prof Vladyslav Vyazovskiy (PI) and Prof Molnár (Co-PI), MRC Project Grant “Orexinergic projections to neocortex: potential role in arousal, stress and anxiety-related disorders”. Molnar (PI) and Mann (Co-PI) (MR/W029073/1), Einstein Stiftung Berlin with Prof Britta Eickholt at Charité-Universitätsmedizin Berlin, Germany as part of being Einstein Fellow at Charité Universitätsmedizin Berlin, Oxford Martin School Grant: repairing the brain with 3D-printed neural tissues (Bayley, Szele, Molnár), and St John’s College Research Centre Grant: “Cellular and molecular interactions between neurons and microglia in normal and altered cerebral cortical development”. Collaborator and sponsor for Dr Auguste Vadisiute JRF at St John’s College, University of Oxford.

## Data availability

The data that support the findings of this study are available from the corresponding authors upon request.

## Author Contributions

Z.M. and A.H.S. conceptualised and designed the study. F.SZ. performed immunohistochemistry experiments, morphometrics analyses, density and laminar distribution analyses, correlation analyses, and imaging, and wrote the manuscript. V.S. performed Vglut1 IHC experiments. F.SZ. wrote the manuscript. Z.M. and A.H.S. supervised the project and secured funding. A.M.S., V.S., and A.H.S. performed pilot experiments.

