## Supplementary materials for "Chronic abolition of evoked vesicle release from layer 5 projection neurons disrupts the laminar distribution of parvalbumin interneurons in the adult cortex"

| **Transgenic**  **strain** | **Mouse**  **Genotype** | **Experimental**  **Cohort** | **Timepoint** | **Nr of brains** | **Nr of sections** | **IHC** |
| --- | --- | --- | --- | --- | --- | --- |
| *Rbp4-Cre;Ai14;Snap25^fl/fl^* | Cre-;Ai14;Snap25^fl/fl^ or  Cre-;Ai14;Snap25^fl/+^ (ctrl) | Developmental | P14 | 3 | 4 | *Parvalbumin*  *Vglut1* |
| *Rbp4-Cre;Ai14;Snap25^fl/fl^* | Cre+;Ai14;Snap25^fl/fl^ (cKO) | Developmental | P14 | 3 | 4 | *Parvalbumin*  *Vglut1* |
| *Rbp4-Cre;Ai14;Snap25^fl/fl^* | Cre-;Ai14;Snap25^fl/fl^ or  Cre-;Ai14;Snap25^fl/+^ (ctrl) | Developmental | P21 | 4 | 4 | *Parvalbumin*  *Vglut1* |
| *Rbp4-Cre;Ai14;Snap25^fl/fl^* | Cre+;Ai14;Snap25^fl/fl^ (cKO) | Developmental | P21 | 4 | 4 | *Parvalbumin*  *Vglut1* |
| *Rbp4-Cre;Ai14;Snap25^fl/fl^* | Cre-;Ai14;Snap25^fl/fl^ or  Cre-;Ai14;Snap25^fl/+^ (ctrl) | Adult | 12 weeks | 5 | 3 | *Parvalbumin*  *Vicia villosa* |
| *Rbp4-Cre;Ai14;Snap25^fl/fl^* | Cre+;Ai14;Snap25^fl/fl^ (cKO) | Adult | 12 weeks | 5 | 3 | *Parvalbumin*  *Vicia villosa* |
| *Both male and female mice were used throughout the experiments.* | | | | | | |

**Supplementary Table 1**. Number of brains and number of sections used per experimental cohorts and genotype to determine the density, laminar distribution, developmental trajectory, and Pearson correlation between PV and VVA neurons. The table also indicates the type of immunohistochemical staining performed at each time point.

| **Transgenic**  **strain** | **Mouse**  **Genotype** | **Experimental**  **Cohort** | **Timepoint** | **Nr of cells** | **Nr of sections** | **Measured**  **features** |
| --- | --- | --- | --- | --- | --- | --- |
| *Rbp4-Cre;Ai14;Snap25^fl/fl^* | Cre-;Ai14;Snap25^fl/fl^ or  Cre-;Ai14;Snap25^fl/+^ (ctrl) | Developmental | P14 | 175 | 4 | *Soma area*  *Perimeter*  *Circularity*  *Feret*  *MinFeret*  *Feret angle*  *Roundness*  *Solidity* |
| *Rbp4-Cre;Ai14;Snap25^fl/fl^* | Cre+;Ai14;Snap25^fl/fl^ (cKO) | Developmental | P14 | 131 | 4 |  |
| *Rbp4-Cre;Ai14;Snap25^fl/fl^* | Cre-;Ai14;Snap25^fl/fl^ or  Cre-;Ai14;Snap25^fl/+^ (ctrl) | Developmental | P21 | 180 | 4 |  |
| *Rbp4-Cre;Ai14;Snap25^fl/fl^* | Cre+;Ai14;Snap25^fl/fl^ (cKO) | Developmental | P21 | 236 | 4 |  |
| *Rbp4-Cre;Ai14;Snap25^fl/fl^* | Cre-;Ai14;Snap25^fl/fl^ or  Cre-;Ai14;Snap25^fl/+^ (ctrl) | Adult | 12 weeks | 168 | 3 |  |
| *Rbp4-Cre;Ai14;Snap25^fl/fl^* | Cre+;Ai14;Snap25^fl/fl^ (cKO) | Adult | 12 weeks | 171 | 3 |  |
| *Both male and female mice were used throughout the experiments.* | | | | | | |

**Supplementary Table 2**. Number of cells and number of sections used to perform morphometric analyses of various soma features of PV-positive neurons in the caudoputamen at each time point. The table shows the soma parameters selected for morphometric measurements.

| **Transgenic**  **strain** | **Mouse**  **Genotype** | **Experimental**  **Cohort** | **Time**  **point** | **Cortical ROIs**  **(Local effect)** | **Subcortical ROIs**  **(Global effect)** | **Morphometrics** | **Pearson correlation** |
| --- | --- | --- | --- | --- | --- | --- | --- |
| *Rbp4-Cre;Ai14;Snap25^fl/fl^* | Cre-;Ai14;Snap25^fl/fl^ or  Cre-;Ai14;Snap25^fl/+^ (ctrl) | Developmental | P14 | Primary motor cortex (M1)  Primary somatosensory cortex (S1) | Caudoputamen (CPu)  Globus pallidus, external segment (GPe)  Lateral posterior nucleus of the thalamus (LP)  Mediodorsal nucleus of the thalamus (MD)  Superior colliculus (SC) | Caudoputamen (CPu) | Primary motor cortex (M1)  Primary somatosensory cortex (S1)  Caudoputamen (CPu) |
| *Rbp4-Cre;Ai14;Snap25^fl/fl^* | Cre+;Ai14;Snap25^fl/fl^ (cKO) | Developmental | P14 |  |  |  |  |
| *Rbp4-Cre;Ai14;Snap25^fl/fl^* | Cre-;Ai14;Snap25^fl/fl^ or  Cre-;Ai14;Snap25^fl/+^ (ctrl) | Developmental | P21 |  |  |  |  |
| *Rbp4-Cre;Ai14;Snap25^fl/fl^* | Cre+;Ai14;Snap25^fl/fl^ (cKO) | Developmental | P21 |  |  |  |  |
| *Rbp4-Cre;Ai14;Snap25^fl/fl^* | Cre-;Ai14;Snap25^fl/fl^ or  Cre-;Ai14;Snap25^fl/+^ (ctrl) | Adult | 12 weeks |  |  |  |  |
| *Rbp4-Cre;Ai14;Snap25^fl/fl^* | Cre+;Ai14;Snap25^fl/fl^ (cKO) | Adult | 12 weeks |  |  |  |  |

**Supplementary Table 3**. Cortical and subcortical regions of interest selected to examine the *local* (location of the cell bodies of Rbp4-Cre+ L5 neurons) and the *global* (projection sites of Rbp4-Cre+ L5 neurons) effects of chronically abolishing regulated synaptic vesicle release from cortical L5 projection neurons. The table also indicates the regions of interest for morphometric analyses of PV neurons and for computing the Pearson correlation between PV and VVA.

| **Morphometrics** | | | | | | | |
| --- | --- | --- | --- | --- | --- | --- | --- |
| ***Parvalbumin*** | | | | | | | |
| Development | | | Adult | Trajectory | | | |
| Parameters | **P14** | **P21** | **Adult** | **P14 - P21 ctrl** | **P14 -P21 cKO** | **P21 - Adult ctrl** | **P21 - Adult cKO** |
| Soma area | ns | * | ns | ns | *** | **** | ** |
| Perimeter | ns | ns | ns | ns | ns | ns | ns |
| Circularity | ns | ns | ns | ns | ns | **** | ** |
| Feret | ns | ns | ns | ns | ns | ns | ns |
| MinFeret | ns | * | ns | ns | *** | **** | ns |
| Feret angle | ns | * | ns | ns | ns | ns | * |
| Roundness | ns | ** | ns | ns | *** | ** | ns |
| Solidity | ns | ns | ns | ns | ns | **** | **** |

**Supplementary Table 4**. Summary of the results of morphometric analyses performed on PV neurons in the caudoputamen to determine the short- and long-term impacts of the chronic abolition of evoked vesicle release from L5 projection neurons on the morphology of PV interneurons.

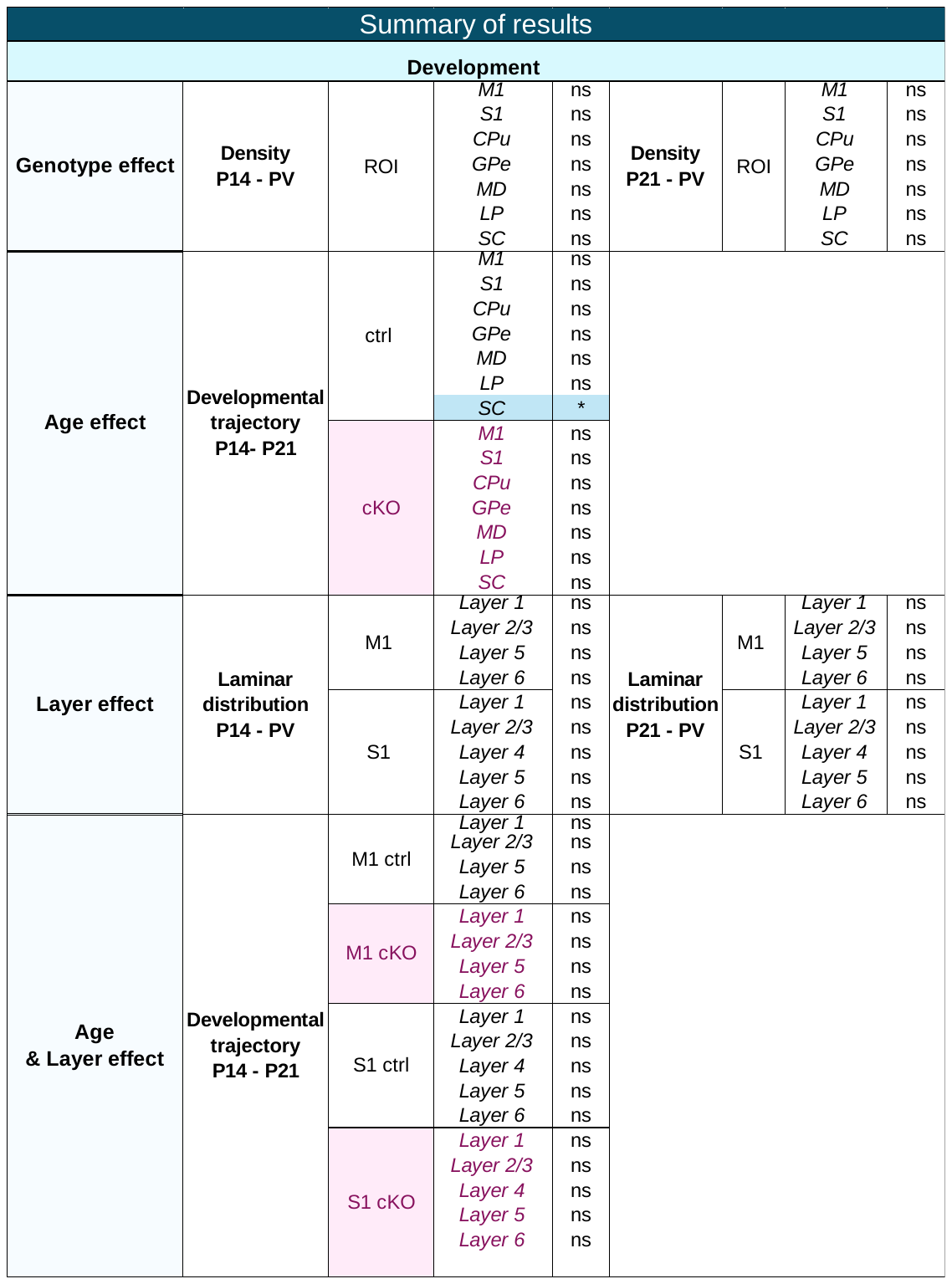

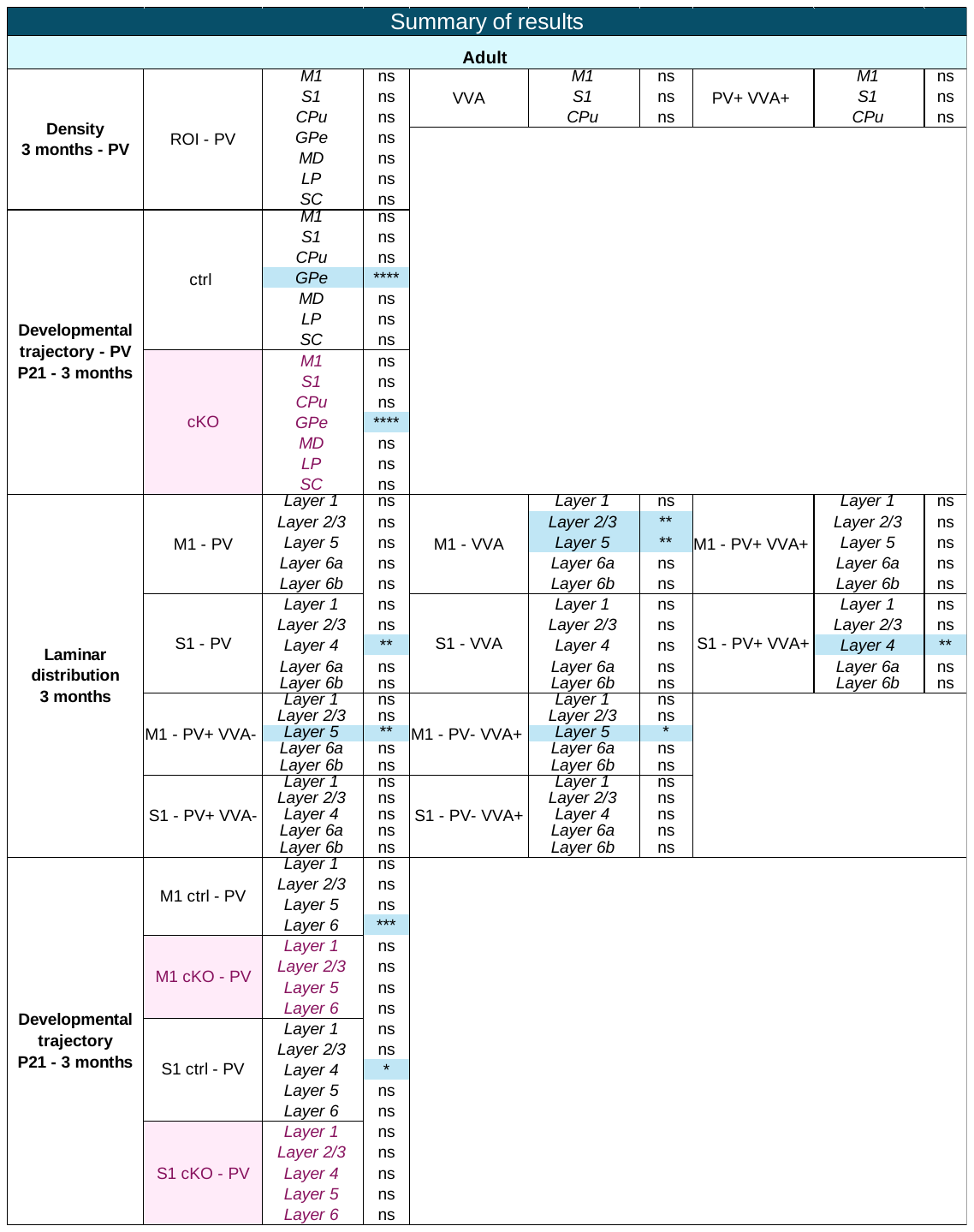

**Supplementary Table 5.** Summary of the results of the density, laminar distribution, and trajectory of PV neurons and their different subpopulations (PV+ VVA+, PV+ VVA-, PV- VVA+) in the cortical and subcortical regions of interest at different postnatal stages and in adulthood.
